# Olive Flowering dependence on winter temperatures - linking empirical results to a dynamic model

**DOI:** 10.1101/2024.02.21.581335

**Authors:** Ilan Smoly, Haim Elbaz, Chaim Engelen, Tahel Wechsler, Gal Elbaz, Giora Ben-Ari, Alon Samach, Tamar Friedlander

**Author notes:** Corresponding Authors: Alon Samach, Tamar Friedlander. Emails: Ilan Smoly, Haim Elbaz, Chaim Engelen, Tahel Wechsler, Gal Elbaz, Giora Ben Ari.

## Abstract

Increasing winter temperatures jeopardize the yield of fruit trees requiring a prolonged and sufficiently cold winter to flower. Assessing the exact risk to different crop varieties is the first step in mitigating the harmful effect of climate change. Since empirically testing the impacts of many temperature scenarios is very time-consuming, quantitative predictive models could be extremely helpful in reducing the number of experiments needed. Here, we focus on olive (*Olea europaea*) – a traditional crop in the Mediterranean basin, a region expected to be severely affected by climatic change. Olive flowering and consequently yield depend on the sufficiency of cold periods and the lack of warm ones during the preceding winter. Yet, a satisfactory quantitative model forecasting its expected flowering under natural temperature conditions is still lacking. Previous models simply summed the number of ‘cold hours’ during winter, as a proxy for flowering, but exhibited only mediocre agreement with empirical flowering values, possibly because they overlooked the order of occurrence of different temperatures.

We empirically tested the effect of different temperature regimes on olive flowering intensity and flowering-gene expression. To predict flowering based on winter temperatures, we constructed a dynamic model, describing the response of a putative flowering factor to the temperature signal. The crucial ingredient in the model is an unstable intermediate, produced and degraded at temperature-dependent rates. Our model accounts not only for the number of cold and warm hours but also for their order. We used sets of empirical flowering and temperature data to fit the model parameters, applying numerical constrained optimization techniques, and successfully validated the model outcomes. Our model more accurately predicts flowering under winters with warm periods yielding low-to-moderate flowering and is more robust compared to previous models.

This model is the first step toward a practical predictive tool, applicable under various temperature conditions.

## Introduction

Flowering in plants requires an initial floral transition, in which a vegetative meristem which previously formed leaf primordia begins to form flower primordia, thus becoming an inflorescence meristem. In annual plants, such as Arabidopsis, at some point in time, a single apical meristem goes through this binary switch to flowering. The time, or the number of leaves formed preceding this switch, is quantitative, depending on the genotype, the environmental conditions, and the plant’s internal state. Once annual plants enter this reproductive stage, they form flowers that develop into seed-containing fruits and eventually die. In woody perennials, typically once a year, in response to an environmental or internal cue, thousands of lateral meristems on a tree can potentially go through a binary switch to flowering. This floral transition can happen either in newly formed vegetative meristems or in previous-year meristems that remain vegetative (Albani & Coupland, 2010). The percentage of meristems that go through this transition can vary between years. In many perennials, a high previous-year fruit load reduces the number of meristems entering the floral transition, in a process termed alternate or biennial bearing (Samach & Smith, 2013). The floral transition in all plants is preceded by initial biochemical changes (in leaves or directly in meristems) that end with a flower-promoting signal reaching the meristem. The FLOWERING LOCUS T (FT) protein acts as this signal, accumulates in leaves, and then moves via the phloem to a nearby meristem where it triggers the expression of genes required for flowering (Corbesier *et al*., 2007). Despite these commonalities, the specific external or internal signals that trigger the accumulation of *FT* transcripts in leaves vary between species (Albani & Coupland, 2010, Andres & Coupland, 2012).

One environmental cue that affects flower induction in some annuals is cold winter temperatures (Kerbler & Wigge, 2023). In several genotypes of Arabidopsis, the transition to flowering is postponed in the absence of prolonged exposure to cold temperatures, namely vernalization (Baurle & Dean, 2006). The expression of several genes required for flowering, among them *FT* is repressed by the MADS-box flowering repressor *FLOWERING LOCUS C* (*FLC*) (Samach *et al*., 2000, Searle *et al*., 2006). With high levels of *FLC* expression at the beginning of winter, *FLC* is later epigenetically silenced during vernalization (Berry & Dean, 2015, Sung & Amasino, 2004). This silencing occurs once a conserved Polycomb Repressive Complex 2 (PRC2) interacts with the PHD protein VERNALIZATION INSENSITIVE 3 (VIN3) at the *FLC* locus. *VIN3* expression is under diurnal regulation with peak levels in the afternoon and very low levels during the night (Hepworth *et al*., 2018). There is a gradual increase in the daily maximum of *VIN3* expression towards the end of winter (Hepworth *et al*., 2018). What causes the slow accumulation of *VIN3* transcripts during weeks of winter? A mechanism was recently suggested in which a protein, NTL8, that upregulates *VIN3* expression is found in high concentration in plants exposed to cold temperatures, and is later diluted during tissue growth, which is promoted under warm temperatures (Zhao *et al*., 2020).

Cold winter temperatures also serve as positive flower inductive cues in several evergreen perennials. Flower induction by cold temperature occurs simultaneously with the accumulation of FT-encoding transcripts in leaves in Avocado (Ziv *et al*., 2014), Citrus (Nishikawa, 2013), Litchi (Ding *et al*., 2015), Mango (Gafni *et al*., 2022, Nakagawa *et al*., 2012), and Olive (Engelen *et al*., 2023, Haberman *et al*., 2017). The floral transition is evident with the formation of microscopic inflorescences towards the end of winter, that later emerge in spring (Samach & Smith, 2013). The molecular events leading to *FT* accumulation during winter in evergreen trees are more of an enigma.

Cold winter temperatures serve as environmental cues in additional plant developmental processes. One important process is the release of buds from endodormancy in deciduous trees. In these species, flower induction occurs during the summer, such that microscopic inflorescences are formed before winter (Goeckeritz & Hollender, 2021). These inflorescences are protected by buds that enter winter dormancy. Three stages of dormancy are described in the literature, with the middle stage, termed endodormancy maintained by the accumulation of Dormancy Associated MADS-box transcription factors (DAMs) or MADS-box proteins similar to SHORT VEGETATIVE PHASE (SVP). Expression of these genes is repressed under cold temperatures, leading to the end of endodormancy (Yang et al., 2021). This repression was suggested by some researchers to be due to epigenetic silencing (Zhu et al., 2020).

Crucially, the number of buds released from dormancy determines the number of emerging inflorescences, which directly affects the tree fruit yield. Following cold winters, the percentage of buds released from dormancy is high, whereas following warmer winters, a high percentage of buds remains dormant in spring. As the release from dormancy requires low temperatures over a prolonged period, the descriptive concept of ‘Chilling Units’ (CUs) or ‘Chilling Hours’ that accumulate throughout winter was introduced, although no molecular accumulation mechanism is known. It was then proposed that a putative ‘chilling requirement’ (CR), namely a minimal number of cold hours accumulated, is a prerequisite for dormancy release. Different deciduous species as well as genotypes/cultivars within a species could vary in their CR (Yang *et al*., 2021). Thus, knowing the specific CRs of different species and varieties enables growers to select genotypes that can provide commercial yields in a specific climatic region. We note though, that even for varieties well-adjusted to their growth region, there is no specific CR that ensures complete release from dormancy of all buds.

The need to study plant response under natural temperature conditions has become critical in the face of recent climatic changes. In particular, recent winters are warmer compared to past winters and if this trend continues, even warmer winters are expected in the coming years (IPCC, 2014). Thus, a cultivar that typically met its CR and provided a high yield in a specific region in previous years, might no longer achieve that in the future (Campoy *et al*., 2011). Moreover, as previous research has already pointed to the inhibitory effect of heat spells on the release from endodormancy, it is concerning that the forecasted increase in such events could compromise yield even if a sufficient cold period is still in place. Yet, the effect of short-term heat spells and the interplay between cold and warm periods were mostly overlooked in previous studies.

A possible strategy to mitigate rising temperatures is picking cultivars with suitable cold requirements per region. However, to accomplish that, a necessary step is a thorough mapping of cultivar temperature sensitivity under realistic temperature profiles and in particular, accounting for the effect of heat spells on flowering. Since the number of temperature profiles for which flowering could be tested empirically is limited for practical reasons, quantitative models could help fill this gap and predict the temperature response under additional profiles. Thus, quantitative models can accelerate breeding programs aimed at alleviating the harsh effects of global warming on crop yields.

Different quantitative models aimed to address this need. The widely used class of ‘summation models’ assumes that trees require a minimal number of CUs for dormancy release. These models simply sum over the number of hours in the temperature range known to promote dormancy release (0 °C to 7.2 °C; Weinberger, 1950). The more sophisticated ‘Utah Model’ (Anderson & Seeley, 1992, Richardson *et al*., 1974) assigns varying weights to different temperatures according to their contribution to dormancy release and specifically negative weights to high temperatures to account for their inhibitory effect. These models are easy to use and have become very popular among farmers. They were applied to various deciduous trees, such as cherry, peach, apple, and pistachio.

Crucially, all summation models only account for the number of hours under each temperature and ignore the order in which these temperatures occurred. Empirical work by Amnon Erez’s group showed that while short heat spells have little effect, long ones strongly suppress endodormancy release, even if the number of cold hours needed is met (Erez et al., 1979a, 1979b). To address this discrepancy with previous models, a different modeling approach was proposed by Fishman et al. (Fishman et al. 1987a, 1987b), called ‘the dynamic model’. This model assumes the accumulation of a hypothetical dormancy-breaking factor (DBF) in response to cold temperatures. In contrast to the summation models, it captures the dynamical process in which the DBF forms, thereby accounting for the sequence of temperatures, rather than the number of hours under each. The dynamic model assumes a 2-step reaction in which the DBF is produced from its precursor, such that both the production and the degradation rates of the precursor are temperature-dependent. The balance between these opposing processes changes with the temperature, such that under low temperatures the DBF accumulates, and under high temperatures, the precursor is degraded, and consequently DBF formation arrests. by this consideration of the dynamics, this model distinguishes between the effects of short and long-term heat pulses. This dynamic model is used to estimate the time in which release from endodormancy in peaches and other deciduous fruit trees is expected.

While the above-mentioned models employed a phenomenological approach, researchers studying the effect of vernalization on flower induction of Arabidopsis attempted to mathematically model the details of the plant temperature sensing mechanisms to make precise predictions regarding the response to different temperature conditions. Antoniou-Kourounioti et al. (Antoniou-Kourounioti *et al*., 2018) proposed an elaborate model for vernalization accounting for *VIN3* and *FLC* levels, as determined empirically. This model includes temperature responses on three time scales: long-term (month), short-term (day), and current (hour). This model too accounts for the sequence of temperatures, similar to the Fishman et al. model and was successfully applied on *Arabidopsis*.

Here we attempt to model the effect of cold winter temperatures on flower induction in the evergreen perennial olive tree. Olive (*Olea europaea*) trees are cultivated primarily for their fruits, which are used for table olives and olive oil. Olive oil is the main source of dietary fat in the Mediterranean diet, which is associated with several health benefits, including the reduction of major cardiovascular events. Annual global consumption of olive oil is about 3.2 million tons, with nearly 95% of this production in regions around the Mediterranean Sea. Olive productivity depends on the number of new nodes formed on one-year-old shoots, the percentage of lateral meristems forming an inflorescence (*i*), and the percentage of inflorescences (Haberman *et al*., 2017) forming and retaining fruit (Wechsler *et al*., 2022). Recently, olive yield has been jeopardized by climatic changes, in particular the elevation of winter temperatures in the Mediterranean region (Ben-Ari *et al.,* 2021; Villalobos *et al.,* 2023). Here we focus on *i* - the percentage of lateral meristems forming an inflorescence – and the environmental factors affecting it. Flower induction in olives during winter requires cold winter temperatures and is not influenced by day length (Haberman *et al*., 2017, Hackett & Hartmann, 1967, Hartmann & Porlingis, 1957). However, the specific cold requirements (CR) differ between olive varieties (Hamze *et al*., 2022, Engelen *et al*., 2023). The expression of two genes that encode proteins similar to FT increases in olive leaves during winter, as well as during controlled conditions of 16/10°C provided in winter or other seasons (Haberman *et al*., 2017). However, if a tree bears a high fruit load in the year before, the accumulation of FT-encoding transcript during winter will be suppressed even under temperatures that otherwise promote flowering, relative to a tree bearing a low fruit load (Haberman *et al*., 2017).

Experiments using potted olive trees of several cultivars subjected to different winter temperature regimes have shown that exposure to temperatures in the range of 5°C - 15.5°C for 13-15 weeks promotes *i*, while warm winter temperatures inhibited flowering (Badr & Hartmann, 1971, Hackett & Hartmann, 1967, Hartmann & Whisler, 1975, Ramos *et al*., 2018, Rubio-Valdes *et al*., 2022). Hence, it was suggested that in olive too, a minimal number of CUs is required to facilitate flowering, but attempts to estimate this minimal number provided confounding results in different olive growth regions. Using a weight function that sums the days matching a different temperature criterion (Denney et al., 1985) yielded a broad range of values: 10-140 in winters of different regions in Texas and 0-200 in different regions of Argentina, Mexico, the USA, and Europe, all of them are regions where olives are grown commercially (Ayerza & Sibbett, 2001, Denney *et al*., 1985). This broad range of values, including zero in one case, suggests this criterion to be too coarse. Others (De Melo-Abreu *et al*., 2004) used a modified ‘Utah model’ to calculate the CRs of different olive varieties.

We have previously collected data on *i* levels in five different olive cultivars and monitored the hourly temperatures throughout nine to eleven winters in several locations in Israel exhibiting a variety of weather conditions (Engelen *et al.,* 2023). Using this data, we attempted to develop a method for calculating CUs that correlated with the level of flower induction observed. We suggested a modified CU accumulation model, assigning Gaussian weights centered around 11 °C to different temperatures. We tested two different variants of the model - Gaussian with positive accumulation only, and Gaussian with both positive and negative accumulation (for high temperatures), and compared to the previous model by De Melo et al. as a reference. We found that all three models were successful in predicting the empirical flowering levels obtained under cold winters yielding high *i*, but much less effective at predicting flowering levels under winters with warm periods in which low to moderate flowering levels were observed (Engelen *et al.,* 2023). We attributed this lack of success under conditions marginal for flowering to the inherent limitation of this class of models, disregarding the dynamic process of temperature change throughout the winter. Rather than trying to further improve the summation models for our purposes, we concluded that a different modeling approach is needed in this case.

In the current work, we employ a multi-faceted approach to study the impact of realistic temperature profiles and specifically heat spells on olive flowering. We began by experimentally testing the effect of heat spells of different durations on olive flowering and gene expression, adding that to our previous database (Engelen *et al.,* 2023). We then proceeded by constructing a new dynamic model that captures the effects of winter temperatures on olive flower induction, inspired by the dynamic model of Fishman et al. for release from dormancy. Our model describes a hypothetical 3-step pathway producing a positive flowering regulator with a precursor that is produced and degraded in temperature-dependent manners. To fit the model parameters, we used 13 sets of measurements of the percentage of flowering meristems *i* in the ‘Barnea’ olive cultivar coupled with hourly temperature data for the entire winter. The parameter fitting in this case is a hard high-dimensional problem, as all parameters should be fitted in parallel, and multiple parameter combinations could yield similar outcomes. To meet this challenge, we constrained the search space using biological considerations and then applied constrained numerical optimization algorithms. We successfully validated the optimization outcomes using the ‘leave-1-out’ procedure. Lastly, we discuss several improvements and extensions of this model and propose future work.

## Materials and Methods

### Exposure to different artificial winter conditions

*2019-2020.* Potted olive ’Barnea’ cultivar trees that were at least 2 years old, were used during the winter of 2019-2020 in the following experiment located at the Faculty of Agriculture of the Hebrew University in Rehovot, Israel. The trees were divided into six treatments, and for each treatment, we used 4-7 trees. In the control treatment, the trees were kept in a net house. Trees from all other treatments were moved on October 24^th^, 2019 to nearby controlled-environment ‘phytotron’ rooms. Here, trees were exposed to day temperatures for 9 hours and night temperatures for 12 hours per calendar day. Changes between day and night temperatures were gradual, spanning 3 h (Sobol *et al*., 2014). During the first ten days, all phytotron-treated trees were exposed to a 16/10 °C regime (Figure 1a). Later on, after exposure to different temperature regimes, all ‘phytotron’- treated trees were moved to a 16/16°C regime, which is neither inductive nor disruptive, for 8-20 days, depending on the treatment (Figure 1a). Then, on the same date, all ‘phytotron’- treated trees were exposed to one week of a 28/22°C regime, to hasten inflorescence elongation. Then all ‘phytotron’- treated trees were moved back to the net-house.

**Figure 1:**
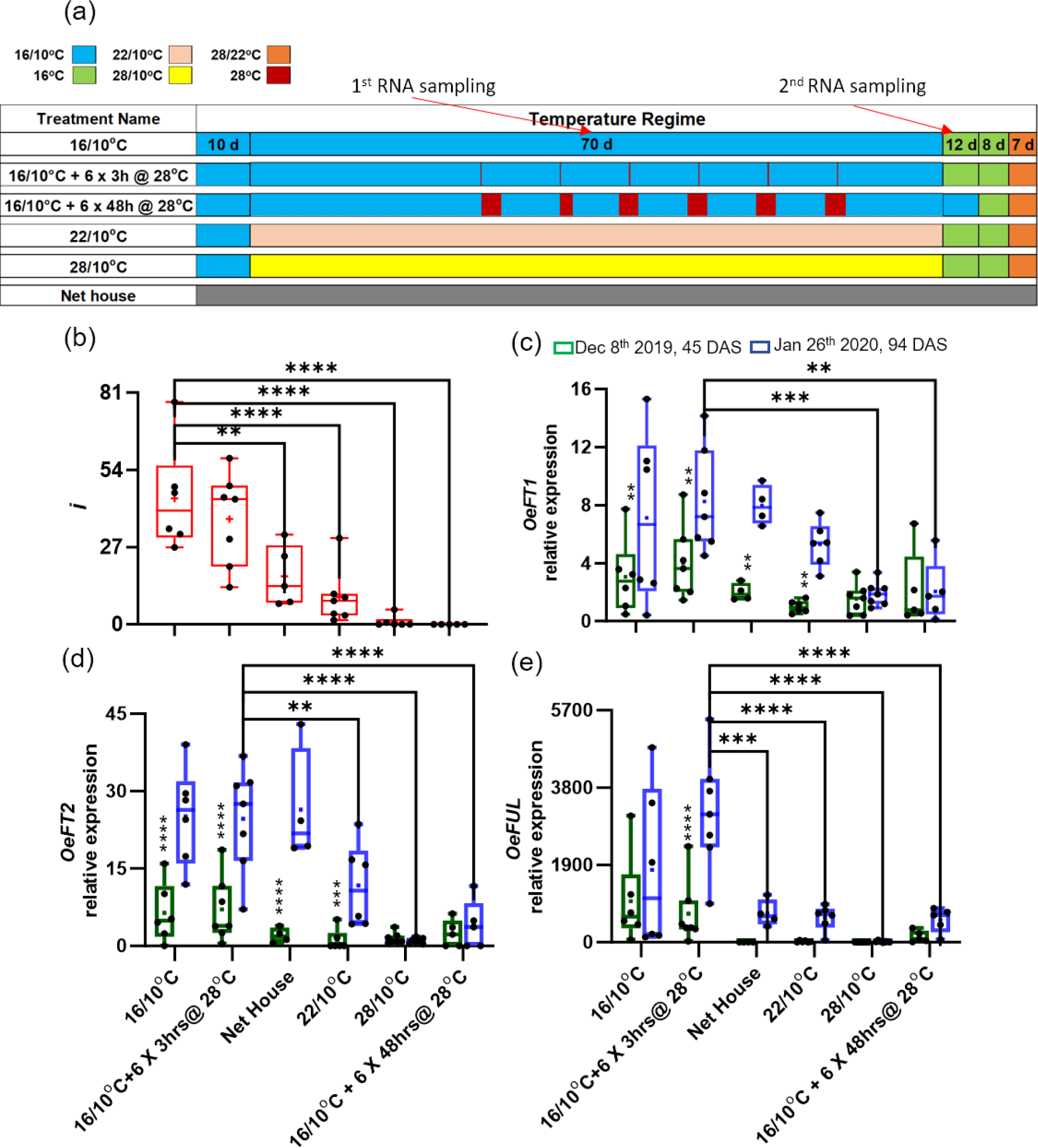
The effect of warm temperatures during artificial winter on flowering and expression of olive flowering time genes. Treatments are different artificial cold conditions during winter compared to outdoors winter conditions (Net house). **(a)** A diagram of the treatments conducted under controlled environments. See the legend for different color codes representing different temperature regimes. Red arrows indicate the two time points in which leaf tissue was collected for RNA extraction and later gene expression. See Methods and Results for details of each treatment. **(b)** The percentage of buds forming inflorescences, *i*. Data is made up of four to seven biological repeats (plants) per treatment. **(c-e)** Relative gene expression in leaves at two sampling dates: December 8th, 2019, 45 days after the treatment start date (green), and January 26th, 2020, 94 days after the treatment start date (blue) of the genes *OeFT1* **(c)**, *OeFT2* **(d)** and *OeFUL* **(e)**. Expression is relative to the OeACT7-1 housekeeping gene. The data is presented as box and whisker plots, the box extends from the 25th to 75th percentiles with the horizontal line within the box at the median, and a “+” at the mean. The whiskers go down to the smallest value and up to the largest, and all data points are presented as black dots. For (a), an ordinary one-way ANOVA Dunnett’s multiple comparison test was performed to study whether the different cold treatments significantly affected tree flowering compared to the 16/10°C treatment. For (c-e), a 2way ANOVA Tukey multiple comparison test was performed to study whether the different cold treatments significantly affected gene expression when compared to the 16/10°C treatment with 3hr pulses of 28°C. In addition, we noted significant increases in gene expression between the first and second sampling dates with asterisks above the earlier sampling date. “*” is p ≤ 0.05, “**” is p ≤ 0.01, “***” is p ≤ 0.001), “****” is p ≤ 0.0001.

In the 16/10°C, 22/10°C, or 28/10°C treatments, the trees were kept under each specific regime for an additional seventy days. After that, the exposure to the 16/16°C regime lasted 20 days. In the short (4hr) heat (28°C) pulse treatment, the trees were also kept in the 16/10°C regime for an additional seventy days, yet within this period they received, once every ∼ 7 days, six 3-hr pulses of 28°C, during the day (Figure 1a). The length of exposure to the following 16/16°C regime was also 20 days. In the long (48hr) heat (28°C) pulse treatment, the trees were kept in the 16/10°C regime for an additional eighty-two days. Within this period, they received once every ∼ 7 days, six 48-hour pulses of 28°C (except for one shorter pulse of 32.5 hrs; Figure 1a). Thus, they were exposed to a similar amount of days under 16/10°C as trees under the 16/10°C regime. The tree’s exposure to the 16/16°C regime was shortened to 8 days (Figure 1a).

### Measurig the Flowering Percent

During the cold exposure period, after the start of natural winter yet before the spring reproductive bud burst had started, branches with flowering potential were chosen at random. Depending on the experiment, 5–10 branches were chosen per plant. Ribbons were tied to mark the previous season’s growth, with the region expected to flower being above, or apically, to the ribbon. After flowering, for each branch, the number of intact buds and the percent of these buds that formed a visible inflorescence (*i*) were recorded. Non-intact buds were those affected by a pest, located in the axil of a pest-ridden leaf or were not in the axil of a leaf at all. *i* values were calculated per branch, with the *i* value for a specific tree being the average *i* value of all recorded branches on that tree. The *i* values of each tree within a treatment were used to calculate the averages and variation within that treatment.

### Tissue Sampling for RNA expression

We collected leaf samples from each one of the trees, for RNA extraction, at two time points during the early morning, 45 (December 8^th^, 2019) and 94 (January 26^th^, 2020) days after the experiment start date (October 24^th^, 2019). Two branches were randomly selected per plant and from each selected branch, two leaf samples were taken. These leaf samples were taken from the beginning of the new growth which is the first node from the opposite end of the apical growth/bud as well as from other different nodes. The leaves from the same tree were combined to represent that tree. The samples were placed immediately in liquid nitrogen and stored in a freezer at -80°C.

### RNA extraction and first-strand cDNA synthesis

See the Materials and Methods section in the supporting information.

### Expression analysis by quantitative real-time PCR

See the Materials and Methods section in the supporting information.

### 2021-2022 28oC heat pulses

See the Materials and Methods section in the supporting information.

### 2021-2022 16/10oC followed by different temperature regimes

See the Materials and Methods section in the supporting information.

### Statistical analysis

Statistical analysis in Figures 1b-e, S1c, and S2c was performed using GraphPad Prism 10.1.2 software. Figure 1b, S1c, and S2c using Brown-Forsythe and Welch ANOVA, followed by DunnettT3 multiple comparisons test. Figures 1c-e, using 2way ANOVA followed by Tukey multiple comparisons test.

### Numerical Optimization Methods used

We used several numerical optimization algorithms to optimize the model parameter values *A*, *B*, *C*, *p*, and *Z*_*scale*_. The optimization minimized the mean square error (MMSE) between the model flowering percent prediction (for given temperatures) and the empirical flowering values. We used constrained optimization to optimize *A*, *B*, *C*, *p*. *Z*_*scale*_ was optimized separately, for each combination of the other four parameters via the Nelder-Mead algorithm (unconstrained optimization). As constraints, we used prior knowledge regarding biologically feasible values. The constraints we used in the optimization were:

- *A* < 0.1.
- 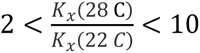

The optimization procedure was run multiple times starting from different initial parameter values of the model, possibly resulting in different estimated parameter values. We compared the final errors achieved using different starting points and chose the model parameter values that yielded the lowest error.

### Local Optimization Algorithms

The local search algorithms were executed using the scipy.optimize.minimize function (Python3).

### Unconstrained optimization

#### Nelder-Mead algorithm (NM)

a derivative-free algorithm that searches the parametric space with a simplex (a convex hull). At each iteration, the algorithm progresses downhill by removing the vertex with the worst cost function value and replacing it with one having a better value. The new point is obtained by attempting to apply reflection, expansion, outside-contraction, and inside-contraction operations (in that order) on the simplex along the line joining the worst vertex with the centroid of the remaining vertices. If this procedure fails (the new candidate points are worst), only the best vertex is retained, and the simplex is shrunk by moving all other vertices toward this value. Formally, given *n* + 1 simplex points 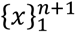 such that *f*(*x*_1_) ≤ *f*(*x*_2_) ≤ ⋯ ≤ *f*(*x*_*n*+1_), the centroid of the top *n* points is 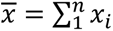 and the new candidate point is obtained by the formula x(*ρ*) = *x̄* + *ρ*(*x*_*n*+1_ − *x̄*). We used the default *ρ* values: 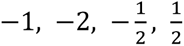 for the reflection, expansion, outside-contraction, and inside-contraction steps, respectively. The shrinking step is done by this formula 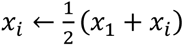. The algorithm stops when both the simplex size (perimeter) and its smoothness (standard deviation in function values across the simplex points) have reached the stop criteria. The predefined parameters we used: initial simplex radius = 0.05, perimeter stop-criteria=0.01, smoothness stop-criteria=0.001.

### Constrained optimization

Interior-point sequential quadratic programming with trust-region (IP) – this algorithm solves at each iteration an equality-constraints sub-problem using quadratic programming, by introducing slack variables (*s*) and solving a sequence of equality-constrained barrier problems for progressively smaller values of the barrier parameter (*μ*):

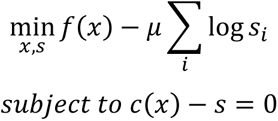

where *c* is a vector of inequality constraints and *s* is a vector of slack variable(s) defined to be positive. This is termed the ‘barrier problem’ which is solved in a two-step procedure: a ‘vertical’ step that minimizes the (new) constraint violation, followed by a ‘horizontal’ step that aims to minimize the (new) cost function requiring that the change in constraint violation is equaled to the one obtained in the ‘vertical’ step. The vertical step is solved with the dogleg method, adapted to allow feasible steps (*s* remains positive, and the trust-region is obeyed) toward the Newton point if it has a lower value (of the quadratic model) than the dogleg step, and the horizontal step is solved with a projected CG algorithm. Three stop-criteria are provided: one for a specific value of *µ*, one for searching across values of *µ,* and one applied to the trust-region radius (which is updated every iteration). The first two stop-conditions are based on the following evaluation function *E*(*x*, *s*, *μ*) = max(‖∇f + ∇*c*λ_c_‖_∞_, ‖sλ_c_ − *μ*‖_∞_, ‖c + s‖_∞_) ‖, where λ_c_ are the Lagrange multipliers for *c.* For simplicity, *µ* represents here a vector of size |*c*| of *µ* values. Note that this is a shortened formula that includes only inequality constraints, as we have in our problem. The full formula is given in Equation 2.3 of (Byrd *et al*., 1999) Both the Jacobian and Hessian are computed at each iteration. The predefined parameters we used were stop-criteria across *µ* =10^−8^, stop-criteria for specific *µ* =0.001, trust-region stop-criteria=0.01, initial trust-region radius=0.05, initial barrier parameter *µ*=0.1, initial constraint penalty = 1.

#### Active-set sequential quadratic programming (SQP)

this algorithm solves at each iteration an (in)equality-constraints sub-problem using quadratic programming applied to the Lagrange function. The algorithm handles the inequality constraints by choosing a working set of active inequality constraints, transforms them into equality constraints, and disregards all the other inequality constraints. The working set is updated according to the step size chosen at the previous iteration. The algorithm stops when it cannot progress anymore, and all Lagrange multipliers for the sub-problem are non-negative (otherwise, it removes the constraint with the lowest Lagrange multiplier from the working set and tries again). Both the Jacobian and Hessian matrices are computed at each iteration.

##### Parameter Scaling

The optimization procedure is executed on the scaled parameters 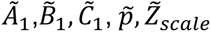, defined such that they all have meaningful values in the range [0,1]:

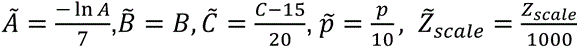

The scaling is crucial for the Nelder-Mead run since both the initial simplex and stop criteria depend on the problem scale. Constrained optimization algorithms are scale-free algorithms since they consider scaling based on the exact Hessian.

##### Initial points

The optimization process was initialized with 192 starting points with different values for the optimized parameters with 4, 3, 4 and 4 starting values for *A*, *B*, *C* and *p*, respectively. The values are 0.0, 0.25, 0.5, 0.75 for *A*, 0.2, 0.45, 0.7 for *B*, 0.1, 0.35, 0.6, 0.849 for *C* and *p*. The optimization ran for 200 iterations from each starting point. This procedure was implemented using scipy.optimize.brute package (Python3). *Z*_*scale*_ was optimized separately, for each combination of the other four parameters via the Nelder-Mead algorithm starting from 5 different initial values: [0.25, 1.1875, 2.125, 3.0625, 4] × *median*_*CU*_, where *median*_*CU*_ is the median value across all treatments of *z*.

##### Hardware

the optimization was run on a local server with dual CPU Intel Xeon 6138, RAM 512 GB, and 40 CPU cores (80 threads).

## Results

### Warm temperatures during winter reduce the flowering level *i*

To better understand the negating effect of warm temperatures during winter on flowering, we exposed small olive trees of the ‘Barnea’ cultivar, grown in pots, to either a natural winter) net house) or to controlled-environment conditions (Figure 1a). Starting October 24^th^, 2019, all controlled-environment treatments were initially exposed for 10.5 days to a 16/10°C daily temperature regime consisting of 16°C day temperature for 9 hrs and 10°C night temperature for 13 hrs, with the remaining 2 hrs a transition period from day to night temperatures (Figure 1a). After this initial term, one group of plants was kept at 16/10°C, another group was moved to a 22/10°C daily temperature regime, and a third group to a 28/10°C daily temperature regime, all for 70 additional days (Fig. 1a).

While *i* levels under 16/10°C were 40%, *i* levels of the trees moved to 22/10°C and 28/10°C were as low as 10% and 1%, respectively (Figure 1b). This result confirms previous studies on the inhibiting effect of warm temperatureExpression levels of s during winter on flower induction (Haberman *et al*., 2017, Hartmann & Whisler, 1975).

To further elucidate the effect of warm temperature intervals during winter on olive flower induction, we subjected some of the trees remaining in the 16/10°C daily temperature regime to six warm temperature (28°C) pulses every 5-8 days. In one treatment each pulse lasted 3hrs, while in the other treatment, the pulse lasted 48hrs. We found that *i* levels in the 3-hr pulse group were not significantly lower than the 16/10°C treated trees with no warm pulse at all (Figure 1b). In contrast, the trees that received 48-hour pulses of 28°C did not flower (Figure 1b).

We then studied the expression in leaves of genes encoding putative flowering regulators. Expression levels of two FT-encoding genes were previously shown to increase in olive leaves during natural winters (Haberman *et al*., 2017, Wechsler *et al*., 2022). We monitored the expression of *OeFT1* and *OeFT2* genes in the leaves of trees exposed to the different treatments, 45 and 94 days from the beginning of the experiment (December 8^th^ and January 26^th^, respectively). In all treatments, except those that caused least flowering (28/10°C and six 48hr heat pulses), both *OeFT1* (Figure 1c) and *OeFT2* (Figure 1d) levels increased significantly in leaf samples collected on January 26^th^ compared to December 8^th^. As a result, when comparing leaf samples collected on January 26^th^, these two low flowering samples had significantly lower levels of *OeFT1* and *OeFT2* expression compared to the trees growing under the 16/10°C regime with six 3 hrs warm (28°C) temperature pulses (Figure 1c-d). In Arabidopsis, the FRUITFULL (FUL) MADS-box transcription factor acts downstream of FT, with levels increasing in both leaves (Teper-Bamnolker & Samach, 2005) and apical meristems (Torti *et al*., 2012). We identified an olive gene encoding a protein similar to FRUITFUL (OE9A106210; *OeFUL*; supplemental text) and monitored its expression in the above leaf samples. In trees growing under the 16/10°C regime with six 3 hrs warm (28°C) temperature pulses, there was a significant increase in *OeFUL* expression in leaf samples collected on January 26^th^ compared to December 8^th^ (Figure 1e). If indeed OeFT proteins are the only ones upregulating *OeFUL* in olive leaves during winter, they are acting during the winter and not only at its end. When comparing leaf samples collected on January 26^th^, the trees exposed to the 16/10°C regime with six 3 hrs warm (28°C) temperature pulses had significantly higher levels of *OeFUL* in leaves compared to trees in the Net house, 22/10°C regime, 28/10°C regime, and the 16/10°C regime with six 48 hrs warm (28°C) temperature pulses.

Examining the relation between *i* levels and gene expression, we observe that the treatment yielding the least flowering (16/10°C regime with six 48-hrs 28°C pulses) had significantly lower levels of expression of all three genes, compared to the treatments with highest flowering (16/10°C regimes with or without six 3-hrs 28°C pulses). Expression of *OeFUL* in leaves was significantly higher in the 16/10°C regime compared to the 28/10 °C regime in both dates. If indeed OeFT proteins are the only ones upregulating *OeFUL* in olive leaves during winter, they are acting throughout the winter and not only at its end.

The stronger effect on expression of the flowering-time-genes, and on *i* of the 48-hrs pulse of 28°C compared to the 3-hr pulse is most likely due to their different durations. Alternatively, the difference in tree response to the two types of pulses may also be due to the different times of the day in which these pulses were applied. Such a time-of-day effect of warm spikes was previously reported for *FLC* silencing in Arabidopsis (Antoniou-Kourounioti *et al*., 2018). To test the possibility for a time-of-day effect in olives, we performed a further experiment comparing between five weekly 3hr 28°C pulses given at either noon or evening (Figure S1). We found no significant difference in *i* between these two treatments suggesting little role for the time of the day at which the pulses were applied.

In our initial experiment, the first heat pulse was provided after 33-days of a 16/10°C temperature regime (Figure 1a). Since trees exposed afterward to several 48-hour pulses at 28°C did not flower (Figure 1b), we conclude that this 33-days cold period was insufficient to secure flowering. We thern asked whether a longer cold period can guarantee flowering, even if high temperature pulses are applied later on. To test that, in the winter of 2021-2022, ‘Barnea’ plants in pots were moved to 16/10°C control conditions from October 25^th^ until December 12^th^ (48 days) and then moved to four different temperature regimes (including regimes with one 4-5hr pulse of temperatures above 25°C) for 21 days (Figure S2a-e). The *i* levels (32-39%) found in these trees were not significantly different between trees subjected to the four different temperature regimes (Figure S2f). Based on these experiments, we conclude that the flowering decision in these trees is made in the time frame between 33 to 48 days following the beginning of the cold period.

### The flowering model

To capture the dependence of *i* values on the cumulative effect of temperatures during the winter months, we constructed a dynamical model, inspired by the dynamical model previously studied for peach dormancy release (Fishman *et al*, 1987a, 1987b) – see Figure 2. An essential component of our model is an unstable intermediate, which we call ‘the temperature-sensitive mediator’ (in the following denoted by *x*). Both the production rate *K*_0_(*T*) and the decay rate *K*_*x*_(*T*) of *x* are temperature-dependent. To ensure a bell-shaped temperature dependence of chilling accumulation, based on previous experiments (Engelen *et al*., 2023, Hartmann & Porlingis, 1957), we defined *K*_0_(*T*) the production rate of *x* to be a Gaussian function with a maximum at the temperature known to be optimal for olive flowering (11 °C) and such that temperatures above 18 °C and below 4 °C have a negligible contribution – see Figure 2b. We defined *K*_*x*_(*T*) the decay rate of *x* to be a sigmoid function, such that for high temperatures the decay rate saturates to its maximal value and for low temperatures, it reduces to a minimal basal decay rate. The exact shape of *K*_*x*_(*T*) depends on three parameters *A*, *B*, *C*, which we later fit using our empirical data (Figure 2c). This definition of the decay rate essentially facilitates the known effect of ‘chilling negation’ by dissipation of the recently accumulated ‘chilling units’ in response to high but not to low temperatures, as found empirically (Fishman *et al*., 1987a). *x* later regulates the production of the ‘intermediate flowering regulator’ (denoted by *y*) which in turn regulates the production of the ‘positive flowering regulator’ *z*. We assume that *y* decays at a constant rate. Analysis of meristem images taken by electron microscope in different developmental stages showed that changes in the meristem could be observed at least two weeks before the inflorescences are observable by the bare eye (Haberman *et al*., 2017). Hence, we decided to collect temperatures only until two weeks prior to the observation of inflorescence emergence by the eye. Yet, in many winters, *y* levels reach their maximum earlier than that last measurement date. Since the exact date on which the flowering decision is made is unknown, we added to the model the stable factor *z*, whose production is regulated by *y*, to avoid sensitivity of our results to the exact end date of the temperature measurement accounted for. *z* represents a hypothetical positive flowering regulator that accumulates throughout the winter season, whereas *x* and *y* form a simplified regulatory pathway of its production and accumulation. We then assert that *z* levels accumulated until floral evocation (two weeks before inflorescence emergence), determine the tree *i* levels. The model is captured by the following scheme (Figure 2). The model parameters and their units are summarized in Table 1.

**Figure 2:**
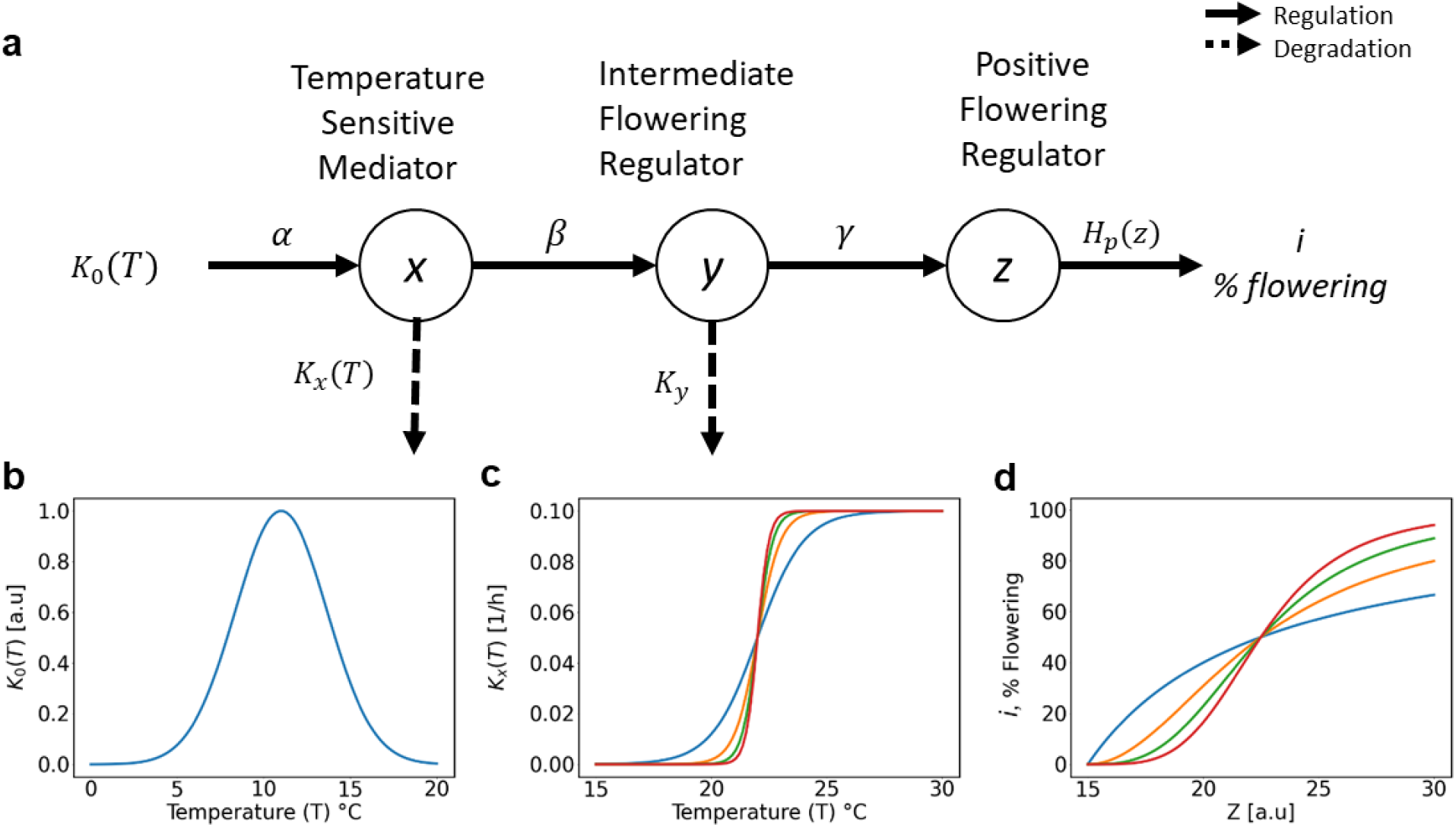
The flowering dynamic model. **(a)** A schematic overview of the proposed dynamic model, which responds to external temperatures throughout the winter and determines the tree flowering level (*i*, % flowering). This hypothetical pathway receives as an input the external temperature *T*, which is then weighted by the function *K*_0_(*T*), here taken as a fixed Gaussian with a maximum at 11_o_C **(b)**. This protein synthesis pathway is composed of three dynamic variables: *x*, *y*, and *z*, which are the levels of the Temperature Sensitive Mediator (TSM), Intermediate Flowering Regulator (IFR), and Positive Flowering Regulator (PFR) respectively. *x* sums over the hours at the desired temperature range given by *K*_0_(*T*), but simultaneously also decays at a temperature-dependent rate *K*_*x*_(*T*) **(c)**, which we define as a logistic function. *x* then gives rise to the production of *y*, that also degrades at a constant rate *K*_*y*_, and *y* in turn leads to the production of *z*. We then convert the *z* levels accumulated by the end of the winter, to the tree flowering level (*i*) using a Hill function **(d)**. We use solid arrows to describe regulation exerted by one variable on another and dashed arrows to designate degradation of variable quantities. Here, the source variables, *x*, *y* and *z* produce α, β, γ units of their outcomes per hour, which by choice of units we set to 1, 5 x 10_-3_, 2 x 10_-3_ respectively (Table 1).

**Table 1:** Parameters in the dynamic model. The reported values are either fixed (f) or optimized (o).

| Variables / parameters | Units | Description | Value (f/o) |
| --- | --- | --- | --- |
| $\alpha$ | 1/hrs | Multiplicative effects on the quantities of the dynamic variables. Define the final units of $z$. | 1 (f) |
| $\beta$ | 1/hrs | Multiplicative effects on the quantities of the dynamic variables. Define the final units of $z$. | $5 \times 10^{-3}$ (f) |
| $\gamma$ | 1/hrs | Multiplicative effects on the quantities of the dynamic variables. Define the final units of $z$. | $2 \times 10^{-3}$ (f) |
| $K_0$ | a.u./hrs | Production rate of $x$- $K_0(T) = e^{-\frac{(T-\mu)^2}{2\sigma^2}}$ | $\mu = 11$ (f)<br>$\sigma = 2.655$ (f) |
| $K_x$ | 1/hrs | The temperature-mediated degradation rate of $x$-<br>$K_x(T) = \frac{A}{1+e^{-B(T-C)}} + D$ | |
| $D$ | 1/hrs | The basal degradation rate of $x$ | $5 \times 10^{-3}$ (f) |
| $K_y$ | 1/hrs | The basal degradation rate of $y$ | $2 \times 10^{-3}$ (f) |
| $A$ | 1/hrs | The maximal value of $K_x$ | 0.1 (o) |
| $B$ | 1/<sup>o</sup>Celsius | The activation energy of $x$'s temperature-mediated degradation | 7.153 (o) |
| $C$ | <sup>o</sup>Celsius | Activation temperature of $x$'s temperature-mediated degradation | 22.306 (o) |
| $p$ | - | Hill steepness coefficient. The ultra-sensitivity in the bud-flowering process. | 2.497 (o) |
| $Z_{scale}$ | $[\alpha\beta\gamma][hr]^{\frac{1}{p}}$ | The value of $z$ which produces 50% flowering ($Z_{scale}$) | $Z_{scale} = 594.575$ (o) |

The model input is given as a sequence of hourly temperature (°C) measurements 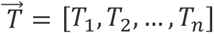. The temperature-sensitive factor *x* is produced at a temperature-dependent rate

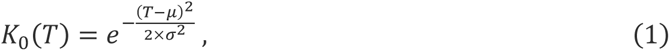

whereas *μ* = 11 °C and *σ* = 2.655 1/°C (Figure 2b). The degradation rate of *x* is a logistic function with an additional constant, representing temperature-independent basal degradation,

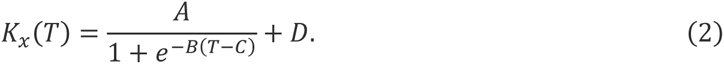

*A* defines the maximal degradation rate of *x*, *B* is the sigmoid slope coefficient which describes the degree of increase in *x*’s degradation in response to a temperature increase and *C* defines the threshold temperature in which *x*’s degradation rate is at half of its maximal value, i.e. 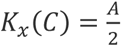 (Figure 2c), and *D* is the basal degradation rate under low temperatures. We later set the values of these parameters using optimization procedures as we detail below. The functions *K*_0_(*T*) and *K*_*x*_(*T*) are updated hourly using the measured temperature *T* (°C) at each time point. The temporal dynamics of *x*, *y*, and *z* are described by the following system of differential equations (3-5):

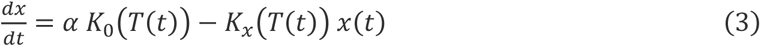

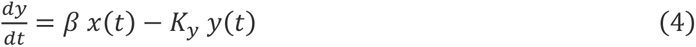

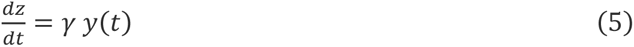

where *T*(*t*) is the temperature at time *t* and *K*_*x*,*y*_(*T*(*t*)) are the degradation rates of *x* and *y* at that time. The parameters α, β, γ have only multiplicative effects on the quantities of the dynamic variables, thus, we fix their values (Table 1) and the final values of *z* are given in units of [*⍺βγ*]. The solution for *x*, *y*, and *z* as a function of the continuous time (*t*) is given by the following equations (6-8):

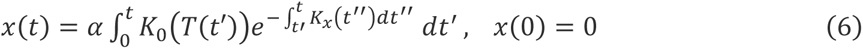

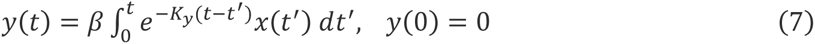

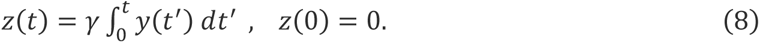

As the temperature data is time-discrete, we use the discrete-time solution of the equations. Then, the values of the dynamic variables can be computed iteratively using the following recursive formulas (9-11):

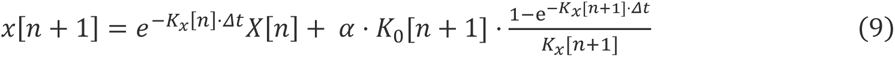

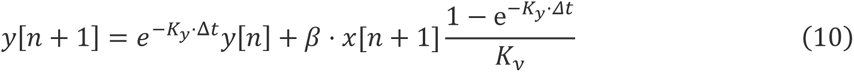

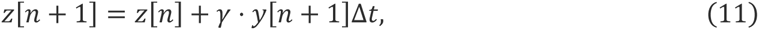

where *K*_0_[*n*] and *K*_*x*_[*n*] are the values of *K*_0_ and *K*_*x*_ (respectively) under the temperature at the *n*-th time point and Δ*t* is the time interval between consecutive temperature measurements – here 1 hour. We define the initial conditions such that, *x*(0) = *y*(0) = *z*(0) = 0, assuming no previous year leftovers of either component.

These equations describe the dynamics of accumulation of the positive flowering regulator *z*, during the winter period. The relation between the regulator level *z* and the tree *i* values is non-linear. Under low levels of the regulator, we assume that flowering is negligible and it saturates to 100 (maximal flowering) under high *z* values. Hence, we chose the Hill function to capture this relation:

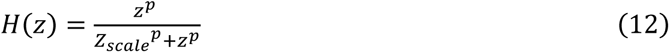

Where *Z*_*scale*_ is a free parameter calibrating the units of *z* such that 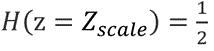. In the following, we refer in the graphs to the scaled variable z/*Z*_*scale*_. *p* is the Hill coefficient that quantifies the steepness of the curve. The Hill equation describes cooperative reactions, where *p* is the cooperativity parameter. Higher cooperativity results in ultrasensitive responses. The Hill function is a convenient choice due to its simplicity and lack of requirement for prior information regarding the details of the modeled pathway (Huang & Ferrell, 1996, Zhang *et al*., 2013). This model implicitly assumes that the rate of the flowering induction process is uniform across all tree branches and all trees. Thus, the output of the Hill equation (Figure 2d; Equation 12) describes the *i* values after a certain temperature regime.

### Dynamics of chilling accumulation in different temperature regimes

We now demonstrate the effects of the model parameters on the chilling accumulation dynamics using simple temperature profiles. In Figure 3 we apply the dynamic model on different temperature cycles in which the night temperature is identical (10°C) and the day temperature is either 16 °C (black), 22 °C (blue), or 28 °C (cyan). The three panels show the temporal dynamics of *x*, *y* and *z* respectively. There is net accumulation of *x* during the ‘night’ (grey) part of the cycle (10°C), and net decay during the warm ’day’ part (white), as both the accumulation *K*_0_ and decay *K*_*x*_ depend on the current temperature. *y* and *z* decay rates are temperature-independent, but since these variables are fed by *x*, the lower *x* is, the lower *y* and *z* are. We demonstrate the effects of different *A* (the maximal degradation rate of *x*) values with either *A* = 0.1 on the left and *A* = 0.01 on the right. A higher decay rate (higher *A*) leads to lower *x* accumulation and hence a more pronounced differentiation between the distinct temperature regimes. Notice that with *A* = 0.1 under the 10/28 °C regime there is no residual *x* left at the end of each warm day.

**Figure 3:**
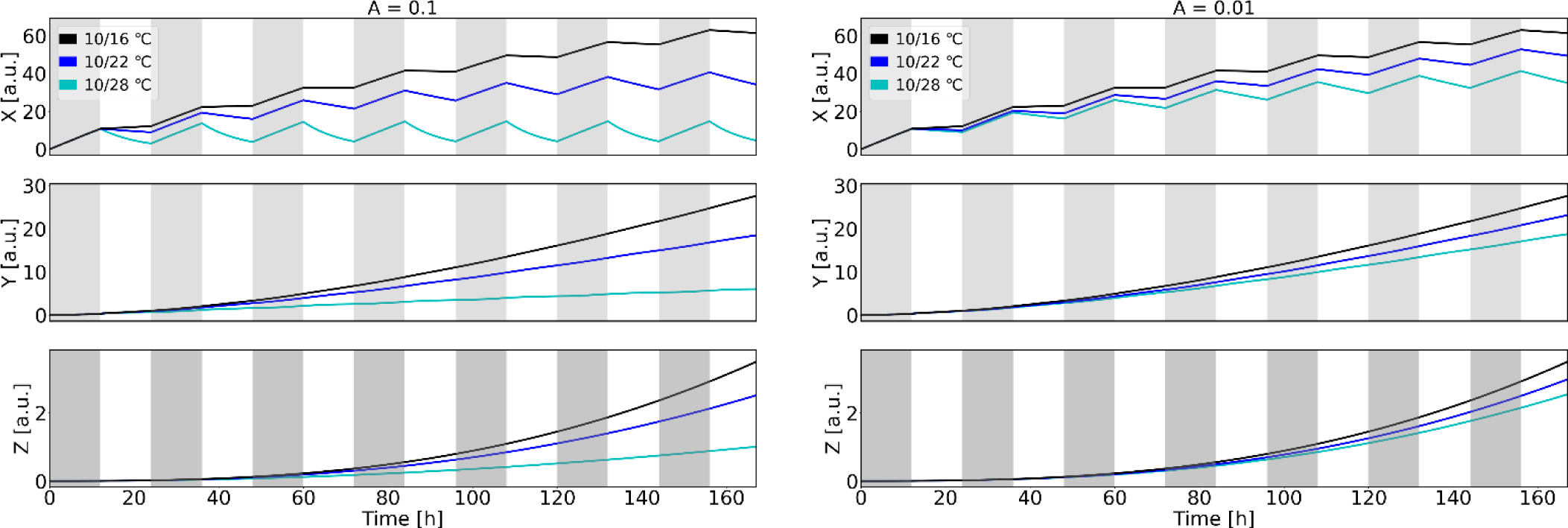
Demonstrating the effect of ***A*** - the maximal degradation rate of ***x***. We demonstrate the dynamics of the model variables *x*, *y* and *z* shown in different rows under a two-temperature cycle mimicking day (12 hrs of high temperatures, white bar) and night (12 hrs of low temperature, grey bar) cycle. In the left column we used *A* = 0.1, whereas in the right column we used *A* = 0.01. Other parameter values are identical in both columns: *B* = 7.153, *C* = 22.306, *D* = 0.005, *K*_*y*_ = 0.002. The different curves indicate different temperature values: 10/16 °C (black), 10/22 °C (blue), and 10/28 °C (cyan). *x* accumulates during low-temperature periods and degrades under high-temperature intervals. While accumulation is identical for all profiles, degradation is more effective under high temperatures. The degradation rate *K*_*x*_ is proportional to *A*, hence the higher *A*, the stronger the degradation effect of high temperatures. Lower levels of *x*, in turn, cause a slower rise of *y* and *z* levels, which in turn can reduce *i* values.

In Figure 4 we examine how warm temperature (28°C) pulses of varying length are manifested in this model. We add to the previous 10/16 °C cycle a single warm temperature (28°C) pulse with a duration of either 3 or 48 hours and demonstrate how *x*, *y* and *z* accumulation is affected, compared to the reference of the 10/16 °C cycle only. We observe that the 48 hrs pulse reduced the accumulated *x* levels to 0.

**Figure 4:**
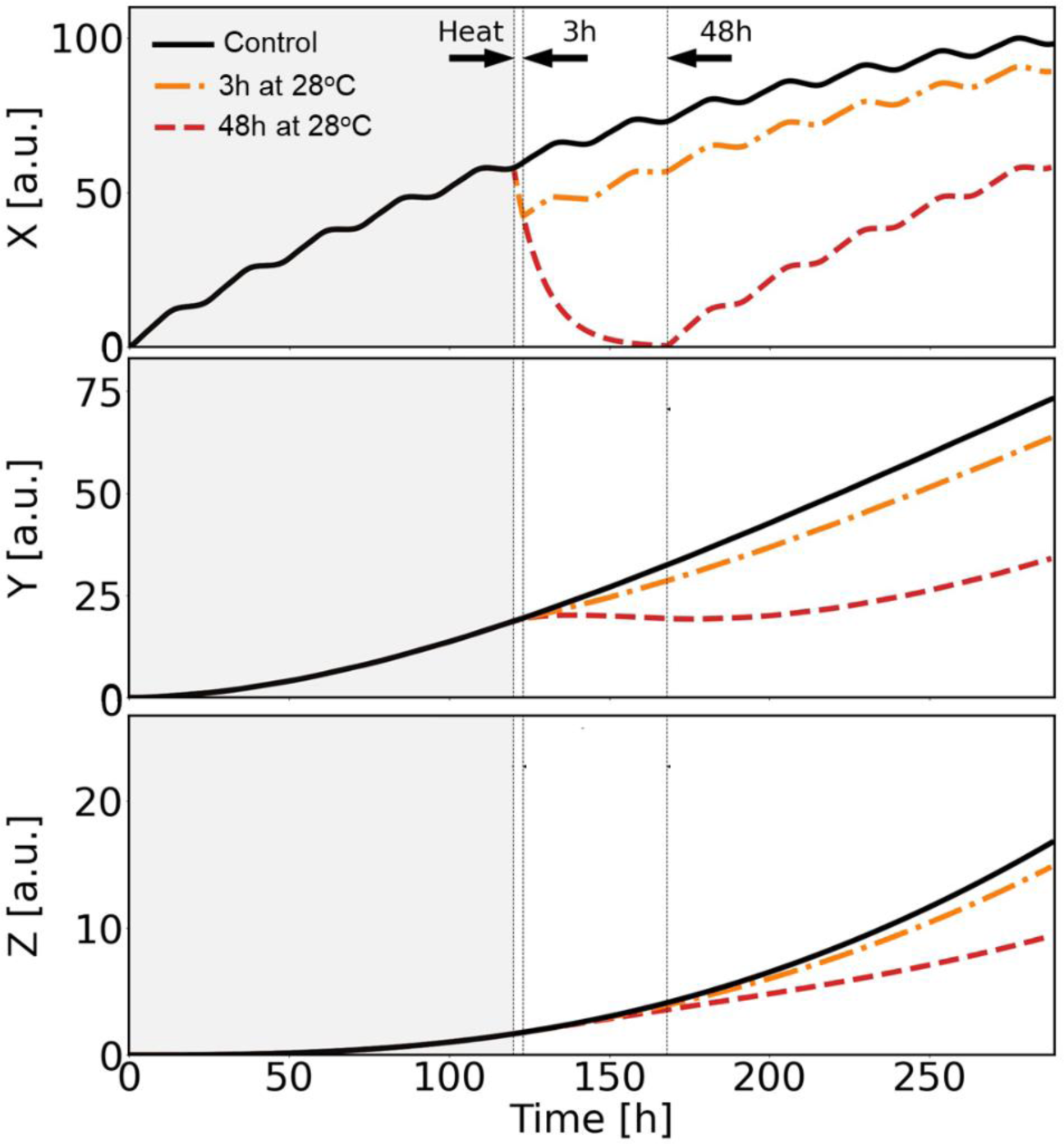
Demonstrating the effect of heat pulses of different lengths on chilling accumulation in the model. We illustrate the accumulation of *x*, *y*, and *z* during a 12-day 10/16°C regime without (black curve) or with a 3 hrs (orange, dash-dot curve) or a 48hrs (red dashed curve) constant 28°C heat pulse embedded once, after 5 days. Gray shades highlight the cold period before the heat event. The 10/16°C daily cycles were smoothed with a sinus filter to mimic natural conditions. We find that a heat period delays chilling accumulation in the model, but if it is short (3 hrs), its effect is minor, whereas a longer (48 hrs) heat period temporarily reduces *x* to zero. Parameter values: *A* = 0.1, *B* = 7.153, *C* = 22.306, *D* = 0.005, *K*_*y*_ = 0.002.

### Fitting the model parameters

The dynamic model of Eqs. 3-5 (continuous time) or Eqs. 9-11 (discrete time) requires the determination of several parameters. We used data from the literature to set the temperature-independent basal decay rates of *x* and *y*. *D* (the basal decay rate of *x*) was set as the median decay rate across 376 proteins in Arabidopsis thaliana leaves during growth, *D* = 5 × 10^−3^ (Li *et al*., 2017). *K*_*y*_ (the decay rate of *y*) value was set as 2 × 10^−3^ which is the decay rate of the 10% most stable proteins in Arabidopsis (Li *et al*., 2017).

To fit the remaining 5 parameter values: *A*, *B*, *C* determining the functional shape of *K*_*x*_(*T*) the decay rate of *x*, and the two Hill function parameters *Z*_*scale*_ and *p* we used the flowering and temperature data we collected. In the fitting procedure (Figure 5) we used as the model input data multiple sets of experiments in which *i* values of several trees were determined in spring and the corresponding temperature profiles throughout the winter months were monitored (Engelen *et al.,* 2023) (see Table S2 for the list of experiments used). To start the optimization procedure, we arbitrarily initialized the model parameter values. At each iteration of the optimization, we calculated the model flowering predictions for all temperature profiles using the current model parameters. We then calculated the model error, defined as the mean square difference between the flowering predictions and the corresponding empirical flowering values, averaged over all experiments. Lastly, we used this error value to update the parameter values to reduce the error further in the next optimization iteration. We repeated these steps multiple times (up to a maximum of 200) until we reached either a predefined error floor or the maximal number of iterations. This entire procedure was repeated multiple times starting from different initial parameter values, each time yielding a (potentially different) set of suggested parameter values for the model. Finally, of all the starting points tried, we picked the parameter combination that yielded the lowest error. The employment of multiple starting points was used to validate the optimization insensitivity to the calculation starting point. The parameter fitting in this case is a hard high-dimensional problem, as all 5 parameters should be fitted in parallel, and multiple parameter combinations could yield similar outcomes. To meet this challenge, we constrained the search space using biological considerations and then applied constrained numerical optimization algorithms. See the Materials and Methods Section for details of the optimization algorithms we used.

**Figure 5:**
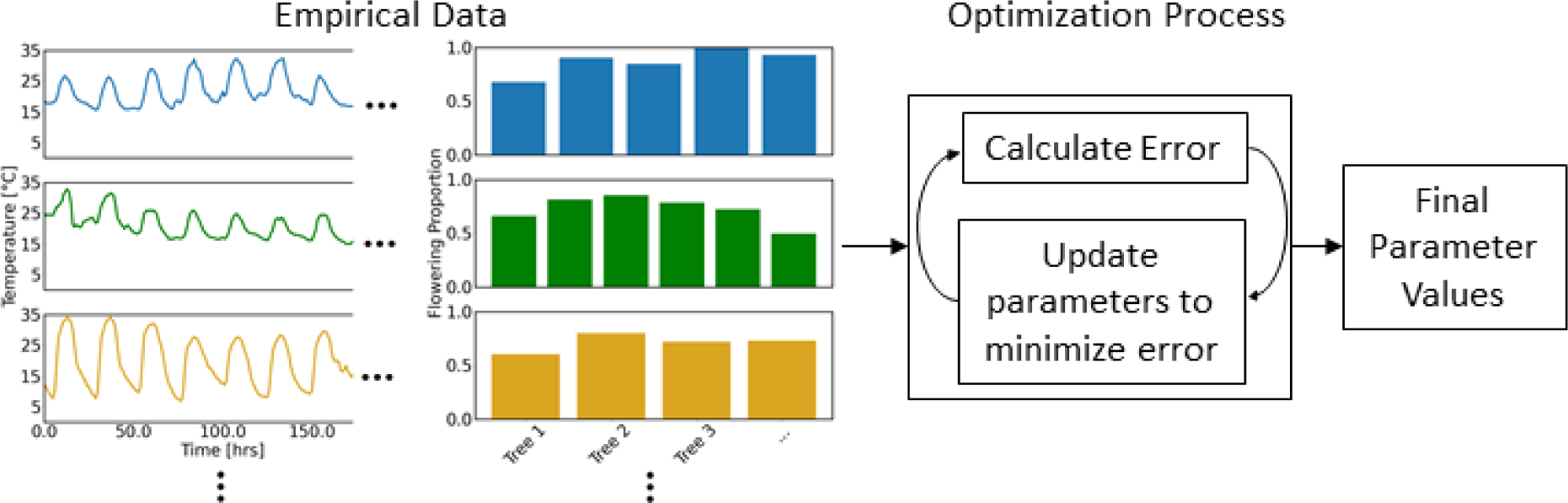
Schematic diagram of the optimization process used for fitting the model parameters. We use sets of empirical olive tree flowering measurements collected in early spring together with the corresponding winter temperature profiles to iteratively optimize the parameters of the dynamic model shown in Figure 2. These include 3 parameters of the temperature-dependent degradation rate *K*_*x*_ and *2* parameters of the Hill function converting *z* to *i*. We begin with multiple arbitrary initial parameter sets. We use these initial parameters to predict flowering under the given temperature profiles, calculate the prediction error, and then use this error to update the parameter values and reduce the error further. We iterate this procedure until either the error or the parameters reach a pre-defined stopping condition. We simultaneously initiate this optimization from multiple starting points, that can potentially provide distinct parameter sets. In most cases however these multiple solutions are similar to each other, demonstrating the insensitivity of the optimization to its starting point.

In Figure 6a we illustrate the optimal Hill function obtained in our parameter optimization (black curve) which maps the *z* values (output of the model regulatory pathway) to *i* values (flowering level). In addition, we illustrate the empirical *i* values of single trees (red dots). The *z* values for each winter were calculated using the optimal parameter values *A*, *B* and *C* by sequentially entering the winter hourly temperatures as the input to the regulatory pathway. Despite the relatively low number of experiments included in the optimization (only 13), we find a good match between the model flowering prediction and the empirical results. In particular, the model successfully predicts cases of very low flowering. Notably, trees under common conditions exhibit considerable variation in flowering levels. One limitation of the model is that it only provides a single number per temperature profile – which we refer to as the mean flowering under that profile – rather than a distribution, as observed empirically.

**Figure 6:**
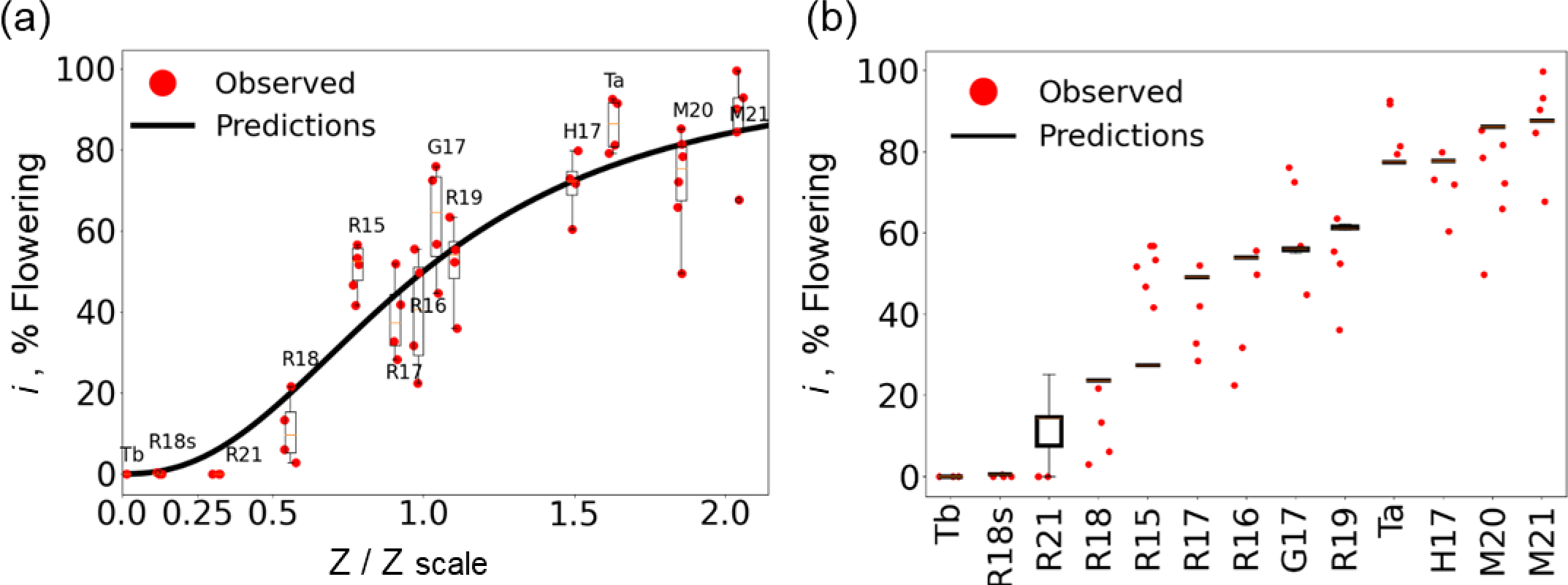
The model flowering prediction and validation. **(a)** The optimal Hill curve obtained maps *i* levels (% flowering) to the regulatory pathway output *z*/*Z*_*scale*_ (black curve, arbitrary units). The red dots illustrate *i* levels measured for each tree and the box plots refer to all trees included in a particular experiment (see Table S2 for experiment short names). We calculated the *z* levels for each experiment by applying the model on the hourly temperature profile of each winter. **(b)**. To validate the model and test its parameter sensitivity to the set of experiments included we applied a “leave-one-out” procedure. Namely, we repeated the optimization multiple times using each time all experiments but one to fit the parameter values. We then used these parameter values to predict flowering for the temperature profile of the single experiment left out. Each column compares the empirical flowering of a single experiment (red dots) to the flowering predicted by the leave-one-out procedure (black box plots). The predictions could potentially span a range of values (as seen for experiment R21), because we ran the optimization from multiple starting points, which could yield different values. Optimal parameter values obtained in the optimization procedure: *A*=0.1, *B*=7.153, *C*=22.306. Fixed parameter values: *D*=0.005, *K*_*y*_=0.002.

### Model validation by a leave-one-out meta-analysis

To test the robustness of our model to the choice of particular temperature profiles we applied the leave-one-out procedure, commonly used in analyses involving many data points. In this procedure, we repeat the parameter-fitting optimization as before, taking each time all the experiments but one. For each excluded experiment we calculate a separate set of the model parameters, obtained independently of that experiment (Table S3). We then apply the model with that parameter set on the temperature profile of the excluded experiment and predict its expected flowering level. Figure 6b shows the leave-one-out results: comparing for each experiment the predicted flowering (black boxes) to the empirical result – each red dot shows a single tree. We find a good match between the predicted and the empirical flowering values, attesting to the insensitivity of the model to the choice of particular experiments, as long as experiments spanning diverse flowering levels are included in the parameter optimization.

We demonstrate the dynamics of the model variables *x*, *y* and *z* under the empirical temperature profiles of Figure 1, comparing chilling accumulation under 16/10 °C, 22/10 °C and 28/10 °C (Figure 7). In the calculation we employ the optimal parameter values, obtained with the different temperature profiles of Figure 6. As expected, we observe the highest chilling accumulation under the 16/10 °C profile, which provides ideal conditions for olive flowering. The 22/10 °C regime leads to a lower *z* accumulation and the 10/28 °C profile exhibits the lowest one, as the high temperatures in these profiles cause chilling negation. This computational result agrees with our empirical data (Figure 1) and bears important practical consequences. Additionally, we calculated the *x*, *y* and *z* dynamics for three natural winter profiles leading to different *i* levels (M21, G17 and R18 - Figure 6a) - see Figure S3 and Table S4 for the calculation details.

**Figure 7:**
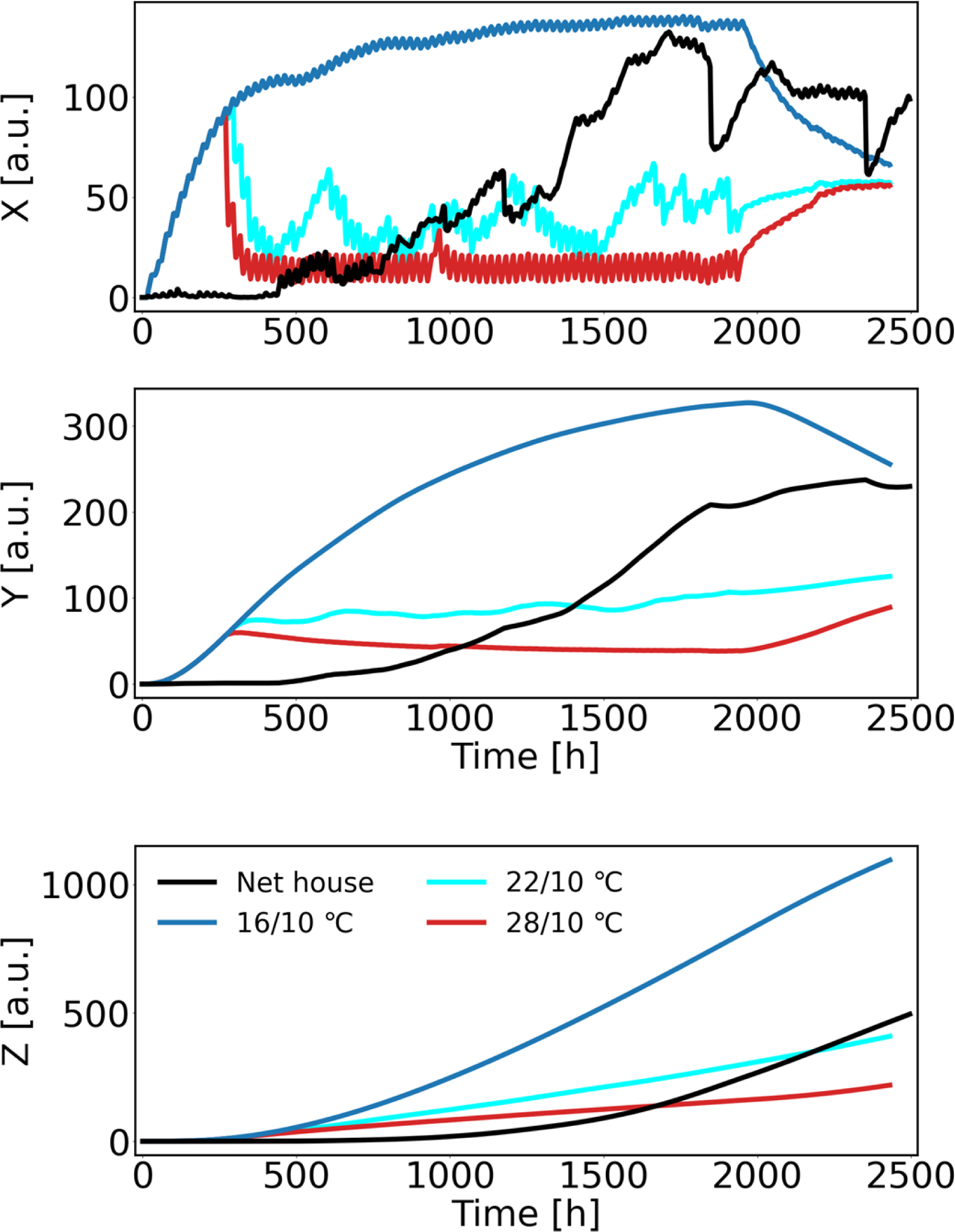
Predicted *x, y* and *z* dynamics for a few of the empirical temperature profiles using the model parameters obtained in Figure 6. The profiles shown here are from the experiment presented in Figure 1: Net house (natural winter in Rehovot 2019-2020) and 3 treatments under controlled temperature conditions of 16/10 °C, 22/10 °C and 28/10 °C. Parameter values used: *A*=0.1, *B*=7.153, *C*=22.3, *D*=0.005, *K*_*y*_=0.002.

### Prediction sensitivity to temperature profile start and end dates

Since *z* cannot decay in our model, end-of-winter warm periods included in the profiles used for optimization and prediction, should have little effect on the predicted *i* levels. Thus, the model is insensitive to the temperature profile end-date. In contrast, it is unclear how to set the earliest date for which temperature should be considered for flowering prediction purposes. Consider also that the beginning of the cold period in the Mediterranean region can vary significantly between years even in the same location. Thus, we tested the sensitivity of our predictions to the starting date of the profile used. In Figure 8 we show the flowering prediction of the model for the different experiments of Figure 6, whereas now we compare the model results for the full temperature profiles to flowering predictions obtained for shortened profiles starting one week later, two weeks later, etc. until 8 weeks later. To avoid bias in the parameter fitting due to the profile shortening, we used for each experiment the model parameters fitted in the leave-one-out procedure in the absence of the particular experiment whose profile was shortened (Table S3). We then used these parameters to predict the flowering levels for the full profile compared to its shorter versions. Impressively, the prediction is insensitive even to profile shortening by 4-5 weeks. Note that our model does not allow for negative chill accumulation, as it only allows for the degradation of already accumulated *x*. In contrast, simple summation models are much more sensitive to the profile starting date, since warm periods at the beginning of the profile cause negative cold accumulation which is deducted from later-occurring cold accumulation. Such a deduction does not exist in our model. This insensitivity to the profile starting date is also exemplified in the *x* accumulation of the natural winter in Figure 7, where there is very little accumulation in the first 500 hours of the profile. Likely, this is also the case in many of the natural winter profiles. Thus, shortening them by a few weeks should have little impact on the flowering prediction.

**Figure 8:**
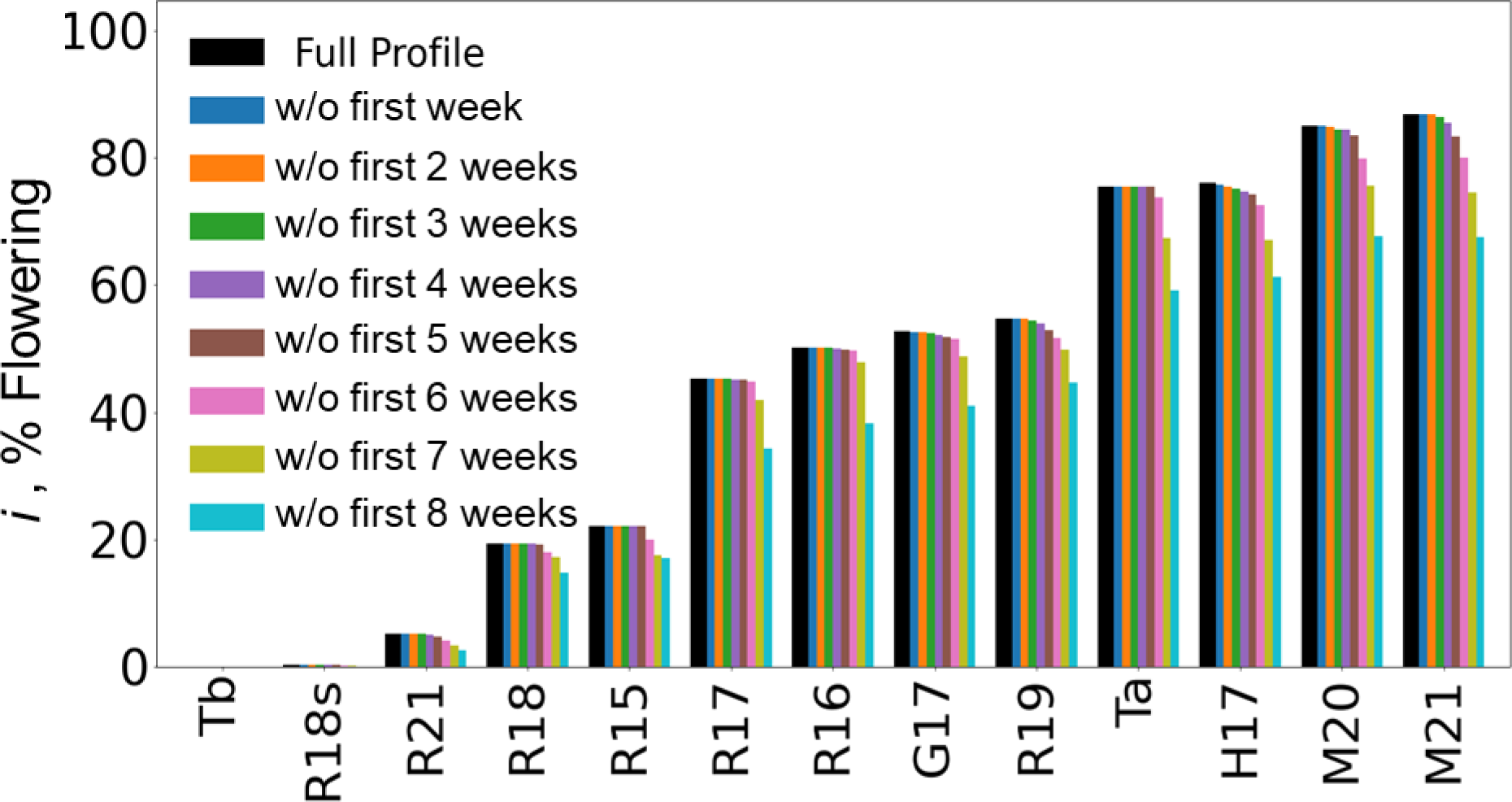
The model flowering prediction is insensitive to the exact starting date of the temperature profile. To test the sensitivity of our flowering predictions to the exact date from which we account for chill accumulation in our model, we used the optimal parameter values obtained in the leave-one-out runs (Table S3) to predict flowering for shortened temperature profiles. For each experiment (name below) we show multiple flowering predictions starting from the full profile (black, as used in Figure 6) and then the profile starting 1 week later, 2 weeks later, etc. until 8 weeks. For all experiments, we find that shortening by up to four-fivre weeks has a negligible effect on the flowering prediction. For all experiments, the full (leftmost bar) profile starts on October 15 and each following bar to the right indicates a subsequent week, with the shortest (rightmost bar) starting 8 weeks later (December 10).

## Discussion

The extent of some developmental switches in plants, for example, the release from bud dormancy in many deciduous trees, and flower induction in several evergreen tree species is determined by the winter temperatures (Kerbler & Wigge, 2023). In this work, we studied the phenomenology of flower induction in response to cold winter temperatures for the ‘Barnea’ olive cultivar, where our main aim was to lay the foundations for a practical tool that could predict olive flowering (and hence yield) based on the temperature profile alone.

### Gene expression and flowering measurement under specific temperature profiles

We began by characterizing the expression in leaves of three genes encoding for proteins known to be involved in the transition to flowering: *OeFT1*, *OeFT2*, and *OeFUL*. We found that the expression levels of all three genes increased over time during the winter cold period. Comparison between different temperature regimes showed that the expression levels of *OeFT2* and *OeFUL* were low and flowering was completely suppressed in trees grown under the 28/10°C regime, whereas both gene expression and flowering were much higher in trees grown under the 16/10 °C regime. We also tested the impact of heat (28°C) spells of different lengths interspersed in an otherwise 16/10 °C temperature regime on the expression of these genes and on the flowering level. Whereas six 3-hour heat intervals had only a minor effect on both gene expression and flowering, six 48hr heat intervals significantly decreased both. We conclude that transcript accumulation during winter of these three flowering genes in the olive tree leaves, is a predictor of its degree of flowering in the following spring and that flowering is not only affected by the number of cold hours accumulated but also by the specific temperature sequence and in particular by the length of uninterrupted heat periods.

These findings are in agreement with previous results showing that transgenic olives overexpressing an FT protein flower all year round (Haberman *et al*., 2017) and that the expression of *OeFT1* and *OeFT2* in leaves of olive trees exposed to different conditions show a positive correlation with *i* levels (Engelen *et al*., 2023, Haberman *et al*., 2017, Wechsler *et al*., 2022). In addition, we identify an olive gene, *OeFUL*, encoding a protein similar to the flower-inducing FRUTIFULL (FUL) from Arabidopsis (Ferrandiz *et al*., 2000; Supporting text)), with a similar expression pattern to the FT-encoding genes in leaves. This is the first report of this gene in olive.

### Model description

Despite the latter findings, the molecular details of temperature sensing, memorizing and processing of this information by the tree remain murky (see below). Thus, to model the dependence of the olive flowering level on winter temperatures, we employed a phenomenological approach. Using the insights gained in the above-mentioned experiments, a basic requirement from such a model was that it should account for the temperature sequence in the entire winter period and in particular for the length of continuous warm and cold periods. This is in contrast to the approach taken by the traditional summation models that only accounted for the number of hours under each temperature and ignored their order. We constructed a model, describing a hypothetical serial regulatory pathway whose input signal is the ambient temperature and whose output is a hypothetical positive flowering regulator *z*, whose levels at the end of the winter directly map into the tree *i* value, namely the percentage of lateral meristems that differentiated into inflorescences. As we found that the timing and expression levels of *OeFT2* and *OeFUL* correlated with cold winter temperatures and subsequent flowering, *z* could be identified with the products of these genes. Still, FT function in the meristems may be inhibited by the levels of other proteins such as TERMINAL FLOWER 1 (TFL1). TFL1 is similar in structure to FT yet with an opposite effect on flowering and can compete with FT (Ahn *et al.,* 2006, Bradley et al., 1997, Kobayashi *et al*., 1999, Lifschitz et al., 2014). The regulation of flowering is significantly more complex than the simplified pathway in our model.

An essential ingredient in this model is an unstable intermediate *x* whose production and degradation rates are both temperature-dependent. We assumed that *x* is produced under cold temperatures, with the highest formation rate at 11°C (Engelen *et al.,* 2023), and decays under high temperatures. We assumed that the *z* levels accumulated until a critical date, in which the flowering decision was made, determined the tree *i* value. Our model uniquely accounts for the tree *i* levels as a continuous variable ranging between 0% (no lateral meristems differentiated into inflorescences) to 100% (all lateral meristems differentiated into inflorescences), in contrast to previous models that considered flowering as an all-or-none phenomenon. Thus, the relation between the regulator level *z* and the tree *i* values should be nonlinear: flowering should be negligible under low *z* levels, and saturate to 100% (maximal flowering) under high *z* values. To capture this nonlinearity, we chose to use the Hill function to describe the relation between *z* and the flowering percentage *i*.

We later used our previously collected *i* values and corresponding winter temperature data (Haberman *et al.,* 2017, Engelen *et al.,* 2023) to fit the model rate parameters by using advanced constrained numerical optimization techniques. We fit the parameters determining the temperature dependence of *x* degradation rate and the Hill function parameters, mapping *z* levels to flowering. Our model uses the entire winter hourly temperature profile as input and sequentially calculates the production and degradation of the flowering regulator until the end of winter. By construction, such a model considers the temporal sequence of temperatures and not only the number of hours under each. Since degradation of the temperature-sensitive intermediate mostly occurs above a threshold temperature, this model exhibits flowering reduction under high temperatures, in agreement with empirical findings. Moreover, as both the production and the degradation of all the pathway constituents are not immediate but paced, only a long warm period could cause the complete depletion of the temperature-sensitive intermediate, which later takes time to rebuild. In contrast, short-term warm intervals will only slightly reduce, but not deplete its reservoir, thus the effect of short-term warm periods on flowering should be mild. This distinction of the model between the impacts of short and long heat periods is in qualitative agreement with our empirical findings showing different responses to 3-hrs and 48-hrs long heat periods.

Given the rate parameters, the model predicts the flowering level expected for a given temperature profile. We validated this model, using the leave-one-out procedure and obtained a good match between the prediction and the empirical results. In particular, the model is more successful compared to previous approaches in predicting flowering intensity under marginal conditions yielding low-to-moderate flowering, where yield is at risk due to global change.

#### Robustness to profile start and end dates

By construction, the model outcome depends on the temperature profile considered as its input, but it is not straightforward how to determine the profile start and end dates. The challenge in determining the profile end date is that it is unknown when exactly the tree reaches its flowering decision. The level of the flowering regulator at this time is thought to determine the tree *i* levels, whereas its accumulation and dissipation beyond that time are unimportant. In the context of our model, if we end the temperature profile too early, we might exclude late-occurring cold periods that could still elevate the flowering regulator level. Conversely, if we end it too late, warm periods at the end could be considered as nullifying formerly accumulated cold periods, while in fact the flowering decision has already been made by that time. To address this uncertainty, we did not include in the model dissipation of the flowering regulator *z*. Thus, if the profile extends into warm periods at the end of the winter, the *z* level already accumulated by that time, is not affected. As the winter start date varies significantly between years in the Mediterranean, it was also unclear when to set the temperature profile start date. To assess this potential error source, we tested the sensitivity of our results to the profile start time. Reassuringly, we found little sensitivity within a range of 4-5 weeks starting from October 15 each year. This desired property of the model is a result of its structure. We assume that all model variables are initialized at zero level, namely that there are no leftover regulators from the previous season. If we initiate the temperature profile earlier than the actual beginning of winter no regulator accumulation, but also no degradation can occur, as long as the temperatures remain high. Both accumulation and dissipation of the flowering regulator only begin when the temperatures become low. This behavior of our model is in stark contrast to that of the summation models that allow for negative cold accumulation, in which initiating the temperature profile at a too-early date might lead to negative net accumulation that would be subtracted from cold accumulation occurring later on. One option to handle this issue is to set the start and end dates, such that the accumulation is maximized (Engelen 2023).

#### Comparison to the dynamic model of Fishman et al

Our model is inspired by an earlier dynamic model constructed by Fishman et al. (Fishman *et al.,* 1987a, 1987b) to account for the dependence of dormancy breaking in deciduous trees on the winter temperatures. This model was the first to account for the ordered sequence of temperatures, rather than the sum of hours under each temperature. As the production and degradation rates in the model followed the Arrhenius law, their values were unlimited, which rendered the Fishman et al. model extremely sensitive to changes in parameter values. In contrast, the rates in our model are bounded. The Fishman model exhibits a dual effect of warm temperatures: both chilling negation and chilling enhancement - the acceleration of anthesis by warm periods at the end of winter. Our model, which aims to predict the level of flowering and not the date of flowering, only shows the former effect.

#### Comparison to summation models

The more prevalent class of models used to predict the long-term temperature effects on developmental processes in trees are summation models (Anderson & Seeley, 1992, De Melo-Abreu *et al*., 2004), that simply sum the number of hours during the winter spent in the desired temperature range, potentially with different weights to different temperatures. The summation models disregard the particular order in which the winter temperatures occurred, thus ignoring the dynamics of accumulation and degradation of relevant factors. In particular, the effect of 48hrs under 28°C during the winter would be the same if given as a single continuous 48hrs pulse or as multiple one-hour pulses every other day. These models might be sensitive to the temperature profile start date, especially if they allow negative cold unit accumulation. Indeed, our previous attempt at using summation models to approximate the flowering level was unsuccessful (Engelen et al., 2023).

The typical lifetime of the unstable intermediate *x* determines the sensitivity of the flowering decision to heat pulses. A very unstable intermediate will respond even to the shortest pulse, whereas the more stable *x* is, the longer the heat pulses it will filter out. The *x* maximal decay rate obtained in our optimization was 0.1 hr^-1^. Comparison to Arabidopsis protein lifetimes (Li *et al*., 2017) shows that this value is at the distribution high end. This could suggest that an active heat-dependent degradation mechanism is at play. In the future, this value can be more accurately determined via controlled experiments testing the response to heat pulses of different lengths.

### Future model extensions

#### Variation in flowering between trees under the same conditions

Our experimental results show considerable phenotypic variation in *i* levels amongst genetically identical trees kept under uniform temperature conditions. Possible explanations for this variation could be additional factors not included in the model that vary between the trees studied, or variation in local concentrations of the flowering regulators between same-tree branches. The current model is deterministic and produces a single flowering level per temperature profile. Future extensions of the model could account for this variation, for example by considering a flowering probability per meristem for each profile and then the model outcome would be a flowering distribution, rather than a single number.

#### Inclusion of previous fruit load

Olives typically have a biennial flowering and yield cycle known as ‘alternate bearing’. We have previously shown that fruit load in one year reduces the flowering level *i* in the following year (Engelen *et al*., 2023, Haberman *et al*., 2017, Wechsler et al., 2022). In the experiments used to estimate the model parameters, we neutralized this previous year effect on flowering by fruitlet removal, mimicking a previous year with no fruit. Hence, the model results should be regarded as the maximal flowering level possible under a specific profile. Previous fruit load is known to reduce flowering, and hence in an actual setting flowering is expected to be lower than the model prediction. Future research should clarify the quantitative relation between the previous year fruit load and the current year flowering and enable the inclusion of this variable too in a more comprehensive version of our flowering prediction model.

#### Additional cultivars

The results reported here apply to the Barnea cultivar. Different cultivars could vary in their cold requirement (Abou-Saaid *et al*., 2022, Aybar *et al*., 2015, Engelen *et al*., 2023, Wechsler *et al*., 2022, Hamze *et al*., 2022), but the mechanistic basis of these different cold requirements remains unclear. One candidate mechanism that could explain this variation might be the level of TFL1 protein in the meristem, which could vary between genotypes. In genotypes with higher TFL1 levels in meristems, the value of *z* which produces 50% flowering (*Z*_*scale*_) would be higher (Haberman *et al.,* 2016). We leave it for future research to explore the mechanisms behind this variation. Future extensions of the model could account for additional cultivars by incorporating different sets of parameter values for different cultivars. Obtaining these parameters will require additional experiments for these cultivars, similar to those used here for ‘Barnea’.

#### Relation to global warming

Olive is a major crop in the Mediterranean. The recent climatic changes, including a gradual increase in mean winter temperatures, aggravated thermal fluctuations, and erratic weather conditions call for studying how its flowering and consequently yield respond to winter temperatures. As winters are getting warmer, cultivars currently receiving sufficient cold to provide adequate flower induction in specific regions might no longer achieve that in the future. Specifically, as previous research has already pointed to the inhibitory effect of heat spells on flowering decisions, it is concerned that the forecasted increase in such events could compromise flowering even if a sufficient cold period is still in place. A possible solution for future rising temperatures could include picking cultivars with suitable cold requirements per region. However, to accomplish that, a necessary step is a thorough mapping of cultivar temperature sensitivity and the construction of accurate quantitative models that will predict flowering under realistic temperature profiles and in particular, account for the effect of heat spells on flowering. Our work provides an important step towards reaching this goal, by the construction of a model that accounts for specific natural temperature profiles and its validation for the Barnea cultivar.

#### Concluding words

More broadly, flowering response to temperature (Kerbler & Wigge, 2023) is an interesting example of a biological response not only to the current signal value, but rather to the cumulative signal level over a long period (Friedlander & Brenner, 2009, 2011). Responses to prolonged signals are also known in animal sensory and neural systems. While the mechanistic implementation varies between systems, their dynamics is surprisingly similar. What mechanisms have trees evolved to sense and collect temperature information? What are the molecular mechanisms that enable a specific temperature to be sensed, recorded and stored for a couple of months, reaching a certain time or threshold when all collected data is analyzed and a decision on the extent of the developmental switch is made? Some new insights into the vernalization response in Arabidopsis suggest one possible mechanism (Jung *et al*., 2020, Zhao *et al*., 2020). Whether flower induction by winter chilling in evergreen trees uses a similar mechanism requires further investigation.

In summary, we report for the first time a phenomenological dynamic model for flowering prediction in olive trees. This work is a first step towards constructing a practical tool that could aid olive growers obtain informed flowering predictions and properly choose cultivars suitable for each region. Our work highlights fundamental biological questions regarding the mechanisms underlying the processing of long-term temperature signals and flowering decision-making, which call for further research.

CUs: ‘Chilling Units’
CRs: Chilling Requirements
FT: FLOWERING LOCUS T

## Acknowledgements

T.F, A.S and G.B-A. acknowledge funding from the Israeli Ministry of Agriculture, grant no. 20-01-0020. T.F. acknowledges funding from Hebrew University and University of Illinois joint research and innovation seed grant program.

The authors declare no conflict of interests.

## SUPPORTING INFORMATION

Additional Supporting Information may be found in the online version of this article at the publisher’s web-site:

**Table S1:** Primers used in this study

**Table S2:** Experimental data used in this study. ‘Type’ indicates whether treatments are based on natural (N) conditions, controlled conditions (C), or both (NC).

**Table S3:** Leave-one-out output parameters.

**Table S4**: Example of *x*, *y*, *z* calculation for a given temperature profile.

### Supporting text

- OeFUL sequence, primers, and alignment
- Additional Materials and methods

## Notes

### Competing Interest Statement

The authors have declared no competing interest.

## References

Abou-Saaid O., El Yaacoubi A., Moukhli A., El Bakkali A., Oulbi S., Delalande M., Farrera I., Kelner J.-J., Lochon-Menseau S., El Modafar C., Zaher H. & Khadari B. (2022) Statistical Approach to Assess Chill and Heat Requirements of Olive Tree Based on Flowering Date and Temperatures Data: Towards Selection of Adapted Cultivars to Global Warming. Agronomy, 12, 2975.

Ahn J.H., Miller D., Winter V.J., Banfield M.J., Lee J.H., Yoo S.Y., Henz S.R., Brady R.L. & Weigel D. (2006) A divergent external loop confers antagonistic activity on floral regulators FT and TFL1. Embo J, 25, 605–614.

Albani M.C. & Coupland G. (2010) Comparative Analysis of Flowering in Annual and Perennial Plants. Current Topics in Developmental Biology. In: Plant Development (ed M.C.P. Timmermans), pp. 323–348. Academic Press.

Anderson J.L. & Seeley S.D. (1992) Modelling Strategy In Pomology: Development Of The Utah Models. Acta Hortic. 313, 297–306

Andres F. & Coupland G. (2012) The genetic basis of flowering responses to seasonal cues. Nature Reviews Genetics, 13, 627–639.

Antoniou-Kourounioti R.L., Hepworth J., Heckmann A., Duncan S., Qüesta J., Rosa S., Säll T., Holm S., Dean C. & Howard M. (2018) Temperature Sensing Is Distributed throughout the Regulatory Network that Controls FLC Epigenetic Silencing in Vernalization. Cell Systems, 7, 643–655.

Aybar V.E., De Melo-Abreu J.P., Searles P.S., Matias A.C., Del Rio C., Caballero J.M. & Rousseaux M.C. (2015) Evaluation of olive flowering at low latitude sites in Argentina using a chilling requirement model. Spanish Journal of Agricultural Research, 13 (1) e0901.

Ayerza R. & Sibbett S.G. (2001) Thermal adaptability of olive (Olea europaea L.) to the Arid Chaco of Argentina. Agriculture, Ecosystems & Environment, 84, 277–285.

Badr S.A. & Hartmann H.T. (1971) Effect of Diurnally Fluctuating Vs Constant Temperatures on Flower Induction and Sex Expression in Olive (Olea-Europaea). Physiologia Plantarum, 24, 40–45.

Baurle I. & Dean C. (2006) The Timing of Developmental Transitions in Plants. Cell, 125, 655–664.

Ben-Ari G., Biton I., Many Y., Namdar D. & Samach A. (2021) Elevated Temperatures Negatively Affect Olive Productive Cycle and Oil Quality. Agronomy, 11, 1492.

Bradley D., Ratcliffe O., Vincent C., Carpenter R. & Coen E. (1997) Inflorescence commitment and architecture in Arabidopsis. Science, 275, 80–83.

Byrd R.H., Hribar M.E. & Nocedal J. (1999) An Interior Point Algorithm for Large-Scale Nonlinear Programming. SIAM Journal on Optimization, 9, 877–900.

Campoy J.A., Ruiz D. & Egea J. (2011) Dormancy in temperate fruit trees in a global warming context: A review. Scientia Horticulturae, 130, 357–372.

Corbesier L., Vincent C., Jang S., Fornara F., Fan Q., Searle I., Giakountis A., Farrona S., Gissot L., Turnbull C. & Coupland G. (2007) FT Protein Movement Contributes to Long-Distance Signaling in Floral Induction of Arabidopsis. Science, 316, 1030–1033.

De Melo-Abreu J.P., Barranco D., Cordeiro A.M., Tous J., Rogado B.M. & Villalobos F.J. (2004) Modelling olive flowering date using chilling for dormancy release and thermal time. Agricultural and Forest Meteorology, 125, 117–127.

Denney J.O., McEachern G.R. & Griffiths J.F. (1985) Modeling the thermal adaptability of the olive (Olea europaea L.) in Texas. Agricultural and Forest Meteorology, 35, 309–327.

Engelen C., Wechsler T., Bakhshian O., Smoly I., Flaks I., Friedlander T., Ben-Ari G. & Samach A. (2023) Studying Parameters Affecting Accumulation of Chilling Units Required for Olive Winter Flower Induction. Plants (Basel), 12. 1714.

Erez A., Couvillon G.A. & Hendershott C.H. (1979a) Quantitative Chilling Enhancement and Negation in Peach Buds by High Temperatures in a Daily Cycle. Journal of the American Society for Horticultural Science, 104, 536–540.

Erez A., Couvillon G.A. & Hendershott C.H. (1979b) The Effect of Cycle Length on Chilling Negation by High Temperatures in Dormant Peach Leaf Buds1. Journal of the American Society for Horticultural Science, 104, 573–576.

Ferrandiz C., Gu Q., Martienssen R. & Yanofsky M.F. (2000) Redundant regulation of meristem identity and plant architecture by FRUITFULL, APETALA1 and CAULIFLOWER. Development, 127, 725–734.

Fishman S., Erez A. & Couvillon G.A. (1987a) The temperature dependence of dormancy breaking in plants: Computer simulation of processes studied under controlled temperatures. Journal of Theoretical Biology, 126, 309–321.

Fishman S., Erez A. & Couvillon G.A. (1987b) The temperature dependence of dormancy breaking in plants: Mathematical analysis of a two-step model involving a cooperative transition. Journal of Theoretical Biology, 124, 473–483.

Friedlander T. & Brenner N. (2009) Adaptive response by state-dependent inactivation. Proceedings of the National Academy of Sciences, 106, 22558–22563.

Friedlander T. & Brenner N. (2011) Adaptive response and enlargement of dynamic range. Mathematical Biosciences and Engineering, 8, 515–528.

Haberman A., Ackerman M., Crane O., Kelner J.J., Costes E. & Samach A. (2016) Different flowering response to various fruit loads in apple cultivars correlates with degree of transcript reaccumulation of a TFL1-encoding gene Plant Journal, 87, 161–173.

Haberman A., Bakhshian O., Cerezo-Medina S., Paltiel J., Adler C., Ben Ari G., Mercado J.A., Pliego-Alfaro F., Lavee S. & Samach A. (2017) A possible role for FT-encoding genes in interpreting environmental and internal cues affecting olive (Olea europaea L.) flower induction. Plant Cell and Environment, 40, 1263–1280.

Hackett W.P. & Hartmann H.T. (1967) The Influence of Temperature on Floral Initiation in the Olive. Physiol Plant, 20, 430–436.

Hamze L.M., Trentacoste E.R., Searles P.S. & Rousseaux M.C. (2022) Spring reproductive and vegetative phenology of olive (Olea europaea L.) cultivars at different air temperatures along a latitudinal-altitudinal gradient in Argentina. Scientia Horticulturae, 304, 111327.

Hartmann H. & Porlingis I. (1957) Effect of Different Amounts of Winter Chilling on Fruitfulness of Several Olive Varieties. Bot. Gaz., 119, 102–104.

Hartmann H.T. & Whisler J.E. (1975) Flower production in olive as influenced by various chilling temperature regimes. J. Amer. soc. Hort. Sci., 100, 670–674.

Huang C.Y. & Ferrell J.E., Jr. (1996) Ultrasensitivity in the mitogen-activated protein kinase cascade. Proc Natl Acad Sci U S A, 93, 10078–10083.

Jung J.H., Barbosa A.D., Hutin S., Kumita J.R., Gao M., Derwort D., Silva C.S., Lai X., Pierre E., Geng F., Kim S.B., Baek S., Zubieta C., Jaeger K.E. & Wigge P.A. (2020) A prion-like domain in ELF3 functions as a thermosensor in Arabidopsis. Nature, 585, 256–260.

Kerbler S.M. & Wigge P.A. (2023) Temperature Sensing in Plants. Annual Review of Plant Biology, 74, 341–366.

Kobayashi Y., Kaya H., Goto K., Iwabuchi M. & Araki T. (1999) A pair of related genes with antagonistic roles in mediating flowering signals. Science, 286, 1960–1962.

Li L., Nelson C.J., Trösch J., Castleden I., Huang S. & Millar A.H. (2017) Protein Degradation Rate in Arabidopsis thaliana Leaf Growth and Development. The Plant Cell, 29, 207–228.

Lifschitz E., Ayre B.G. & Eshed Y. (2014) Florigen and anti-florigen - a systemic mechanism for coordinating growth and termination in flowering plants. Frontiers in Plant Science, 5, 465.

Nocedal J. & Wright S.J. (1999) Numerical Optimization. Springer, New York, USA.

Ramos A., Rapoport H.F., Cabello D. & Rallo L. (2018) Chilling accumulation, dormancy release temperature, and the role of leaves in olive reproductive budburst: Evaluation using shoot explants. Scientia Horticulturae, 231, 241–252.

Richardson E.A., Seeley S.D. & Walker D.R. (1974) A model for estimating the completion of rest for ‘Redhaven’ and ‘Elberta’ peach trees. Hortscience, 9, 331–332.

Rubio-Valdés G., Cabello D., Rapoport H.F. & Rallo L. (2022) Olive Bud Dormancy Release Dynamics and Validation of Using Cuttings to Determine Chilling Requirement. Plants, 11, 3461.

Sobol S., Chayut N., Nave N., Kafle D., Hegele M., Kaminetsky R., Wunsche J.N. & Samach A. (2014) Genetic variation in yield under hot ambient temperatures spotlights a role for cytokinin in protection of developing floral primordia. Plant Cell and Environment, 37, 643–657.

Teper-Bamnolker P. & Samach A. (2005) The flowering integrator FT regulates SEPALLATA3 and FRUITFULL accumulation in Arabidopsis leaves. Plant Cell, 17, 2661–2675.

Torti S., Fornara F., Vincent C., Andrés F., Nordström K., Göbel U., Knoll D., Schoof H. & Coupland G. (2012) Analysis of the Arabidopsis Shoot Meristem Transcriptome during Floral Transition Identifies Distinct Regulatory Patterns and a Leucine-Rich Repeat Protein That Promotes Flowering. Plant Cell, 24, 444–462.

Villalobos F.J., López-Bernal Á., García-Tejera O. & Testi L. (2023) Is olive crop modeling ready to assess the impacts of global change? Frontiers in Plant Science, 14.:1249793. doi: 10.3389/fpls.2023.1249793

Wechsler T., Bakhshian O., Engelen C., Dag A., Ben-Ari G. & Samach A. (2022) Determining Reproductive Parameters, which Contribute to Variation in Yield of Olive Trees from Different Cultivars, Irrigation Regimes, Age and Location. Plants, 11, 2414.

Weinberger J.H. (1950) Chilling requirements of peach varieties. Proceedings of the American Society of Horticultural Sciences 56, 122–128.

Yang Q., Gao Y., Wu X., Moriguchi T., Bai S. & Teng Y. (2021) Bud endodormancy in deciduous fruit trees: advances and prospects. Horticulture Research, 8.

Zhang Q., Bhattacharya S. & Andersen M.E. (2013) Ultrasensitive response motifs: basic amplifiers in molecular signalling networks. Open Biology, 3, 130031.

Zhao Y., Antoniou-Kourounioti R.L., Calder G., Dean C. & Howard M. (2020) Temperature-dependent growth contributes to long-term cold sensing. Nature, 583, 825–829.

Zhu H., Chen P.-Y., Zhong S., Dardick C., Callahan A., An Y.-Q., van Knocker S., Yang Y., Zhong G.-Y., Abbott A. & Liu Z. (2020) Thermal-responsive genetic and epigenetic regulation of DAM cluster controlling dormancy and chilling requirement in peach floral buds. Horticulture Research, 7, 114.

