## supplementary figures for "Olive Flowering dependence on winter temperatures - linking empirical results to a dynamic model"

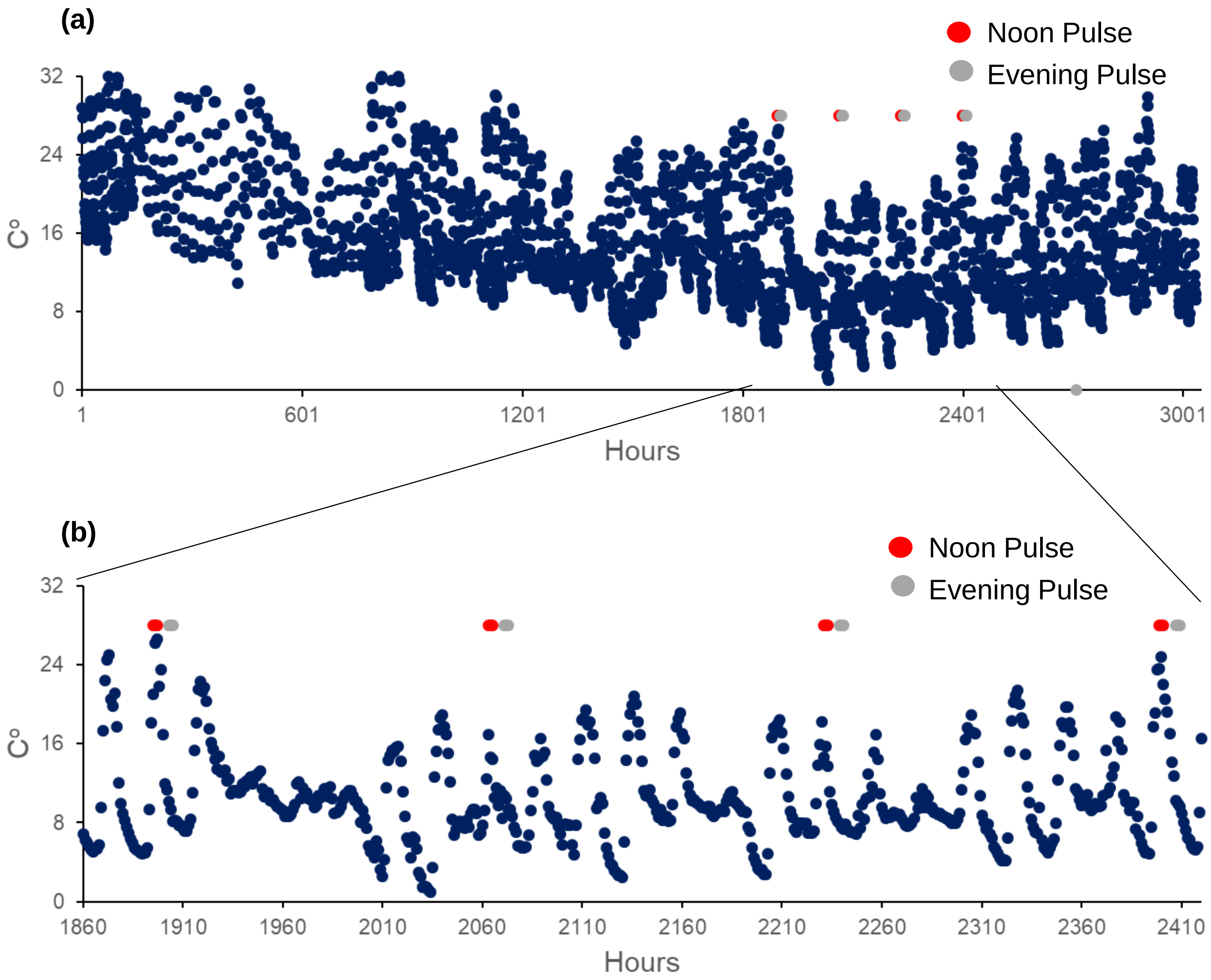


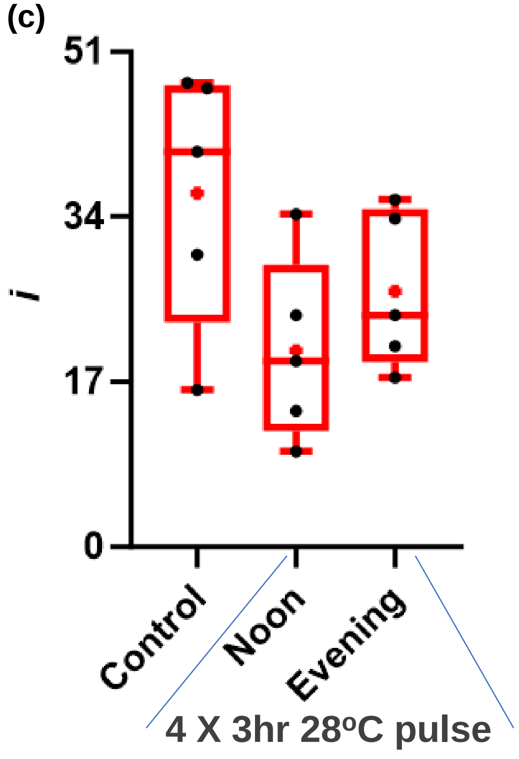


Figure S1 The effect of short heat pulses at different times of the day on olive flowering. (a). Hourly temperatures during the winter of 2021-2022 in Rehovot Israel, from October 25th 2021 until February 28th 2022 (3035 hours). Red dots are the four noon 3hr pulses of 28oC and the grey dots are the four evening 3hr pulses of 28oC given in the two treatments. (b). Insert taken from (a) showing the period in which heat pulses were provided. (c). Flowering percentage (*i*) by treatment. Each treatment contained five 'Barnea' trees (biological repeats). The data is presented using GraphPad Prism 10.1.2 software, the box extends from the 25th to 75th percentiles with the horizontal line within the box at the median, and a "+" at the mean. The whiskers go down to the smallest value and up to the largest and all data points are presented as black dots. No significant differences between treatments were detected.

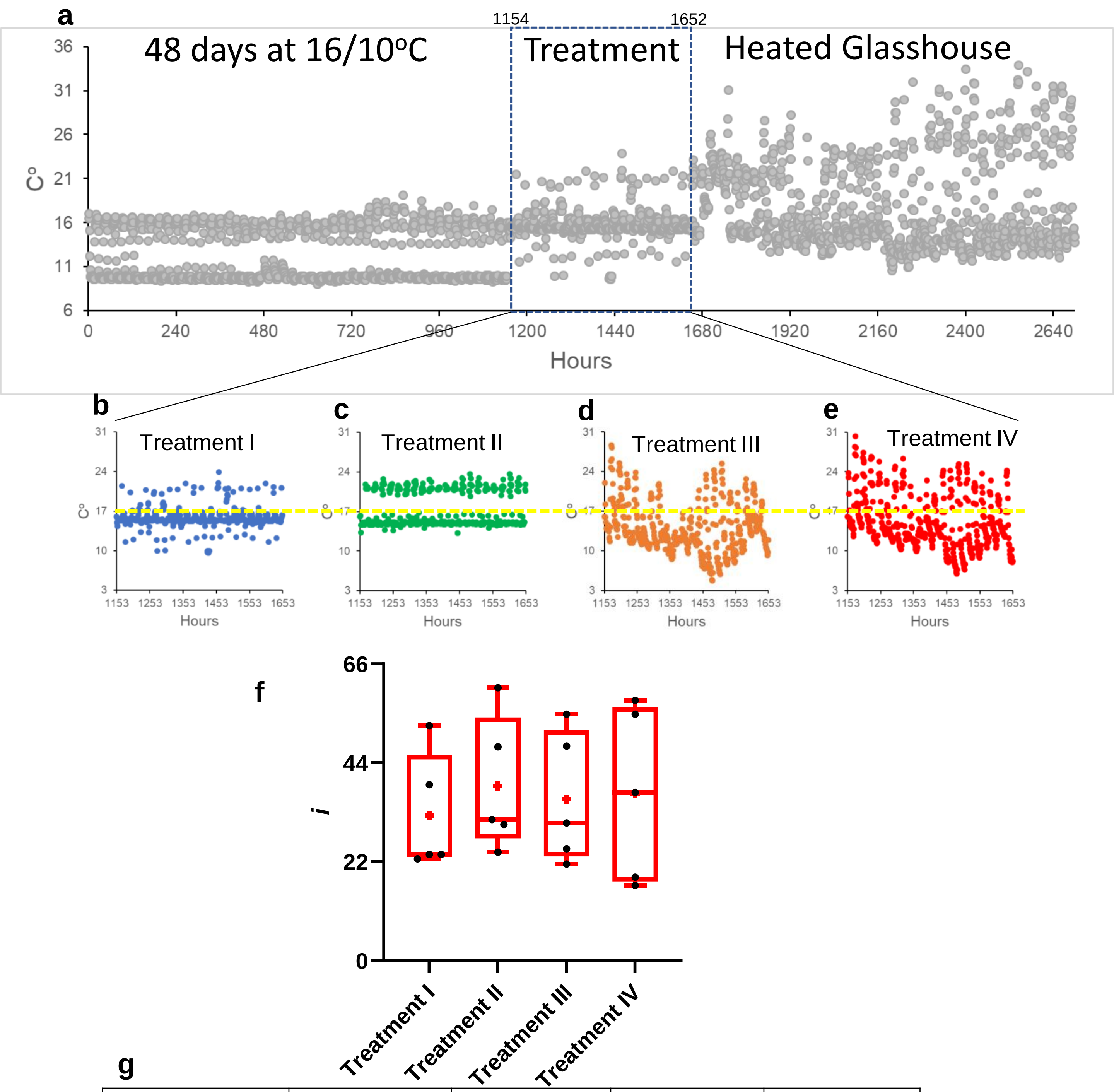


g

| | Treatment I | Treatment II | Treatment III | Treatment IV |
|---|---|---|---|---|
| Pulses ≥ 28°C | 0 | 0 | 1 (2hrs) | 1(3hrs) |
| Pulses ≥ 25°C | 0 | 0 | 1 (4hrs) | 1( 5hrs) |

Figure S2 The effect of different temperature regimes following a 48 day 16/10oC regime on olive flowering. (a). Diagram of hourly temperatures during the whole winter period. All trees received 48 days of 16/10oC followed by 21 days of different temperature regimes (treatments), described in (b-e). Then all trees were kept in the same heated glasshouse until flowering. (f). Flowering percentage (i) by treatment. Each treatment contained five 'Barnea' trees (biological repeats). The data is presented using GraphPad Prism 10.1.2 software, using a box and whisker plots, the box extends from the 25th to 75th percentiles with the horizontal line within the box at the median, and a "+" at the mean. The whiskers go down to the smallest value and up to the largest and all data points are presented as black dots. No significant differences between treatments were detected.

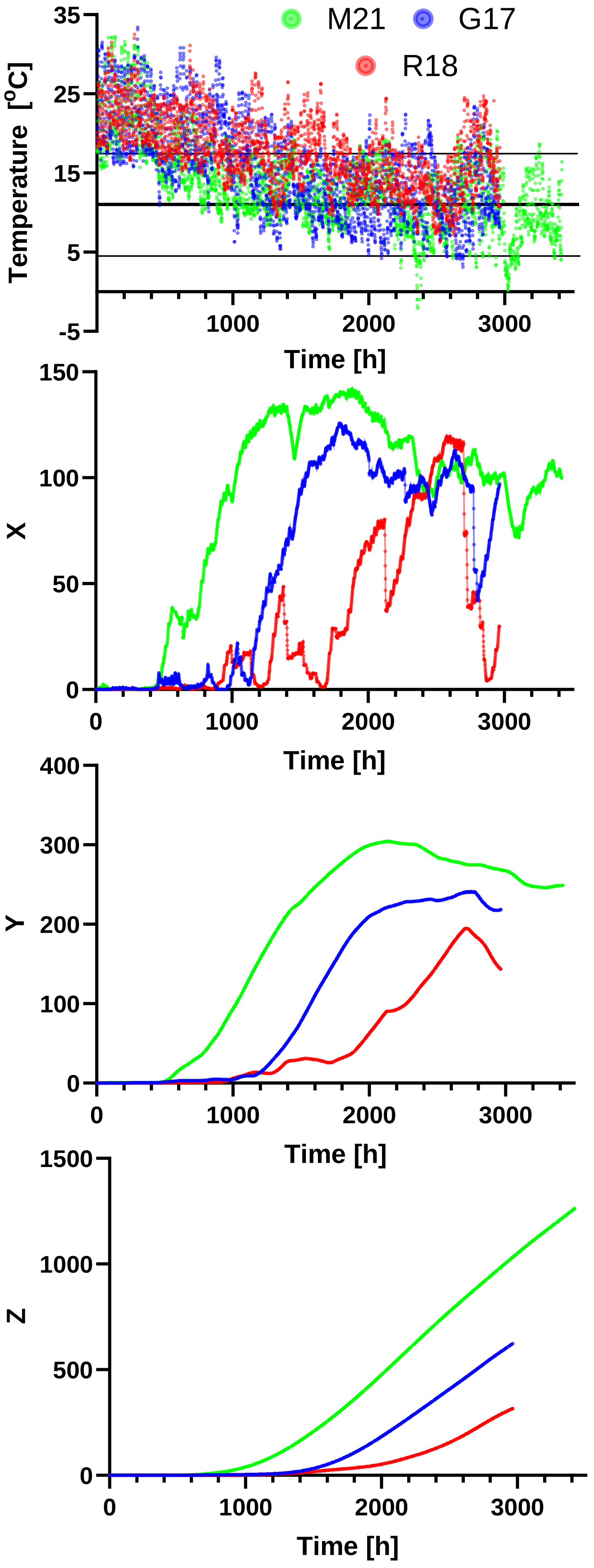


**Figure S3** Predicted $x$, $y$ and $z$ dynamics for three natural winter temperature profiles using the model parameters found in the optimization. The profiles shown are natural winter in Rehovot 2017-2018, Gilat 2016-2017, and Matityahu 2020-2021 (a). Lines in (a) represent optimal temperature (11°C) in bold, as well as upper (17.5°C) and lower (4.5°C) limits of 95% of the fixed Gaussian (Figure 2b). Parameter values used: $A$=0.1, $B$=0.64, $C$=22.3, D=0.005, $K_y$=0.002.
