## supplementary text for "Olive Flowering dependence on winter temperatures - linking empirical results to a dynamic model"

### <u>Information on the *OeFUL* gene and putative protein</u>

>OE9A106210P1

ATGGGAAGGGGAAGAGTTGAGTTGAAGAGAATAGAGAAGAAGATAAATCGGCATGTGACATTTTC
GAAAAGGAGATCTGGGCTTCTCAAGAAAGCTCATGAGATTTCTGTTCTTTGTGATGCAGATGTGG
GTTTGATTGTGTTTTCGCCCAAAGGGAAGCTCTATGAATATGCTACTGGTGCCTGCATGGGGAGG
ATCCTCGAACGATATGAAAGACATTCTTATGAAGAATCACAGCTTACAGCAACGAACATTGAATC
CCCAGTAAGTTGGACGGTAGAATATGCAAAGCTCAAGGCCAGATTGGAGAATTTGCAAACAAGCC
AAAGGCATTACATGGGTGAGGACCTGAATACCCTATGTCTCAAAGAGCTGCAAAATTTGGAACAT
CAGCTTGCTGCTTCTCTTAAACGCATTAGGACTCGCGAGAACCAACTCATGAACAAATCAATTGC
TGAGCTTAAGAAAAAGGACAAGGCATTGCAAGATGAAAACACCTTACTTTTGAAGAAGATCAAAG
AGGAGAAAGAATTATCCCAGAAGTTACAACAGGAACAAAATCATGACATCATATCTTCTTCTGTT
CCACAACTTTTGAGCTCGGGTGAACATGGAGAAGTTGAAGGCCAGTCTCAATCATCTAATGCAGT
AATACCTCCATGGATGATCAACCATCTGCATGAATCCAATAGGAGGGGATAA

Yellow marker: Exon-Exon boders

<u>Green underline</u>: Forward RTPCR Primer

<u>Red Underline:</u> Reverse RTPCR Primer

Translated:

> - 234 codons

MGRGRVELNRIEKKINRHVTFSKRRSGLLKKAHEISVLCDADVGLIVFSPKGKLYEYAT
GACMGRILERYERHSYEESQLTATNIESPVSWTVEYAKLKARLENLQTSQRHYMGEDLN
TLCLKELQNLEHQLAASLKRIRTRENQLMNKSIAELKKKDKALQDENTLLLKKIKEEKE
LSQKLQQEQNHDIISSSVPQLLSSGEHGEVEGQSQSSNAVIPPWMINHLHESNRRG

Alignment with Arabidopsis Protein

```
OeFUL            1 MGRGRVELNRIEKKINRHVTFSKRRSGLLKKAHEISVLCDADVGLIVFSP     50
                   ||||||:|.|||.||||.|||||||||||||||||||||||:|.|||||.
AtFUL            1 MGRGRVQLKRIENKINRQVTFSKRRSGLLKKAHEISVLCDAEVALIVFSS     50

OeFUL           51 KGKLYEYATGACMGRILERYERHSYEESQLTATNIESPVSWTVEYAKLKA    100
                   ||||:||:|.:||.||||||:|:.|.:.||...::....:|.:|:|||||
AtFUL           51 KGKLFEYSTDSCMERILERYDRYLYSDKQLVGRDVSQSENWVLEHAKLKA    100

OeFUL          101 RLENLQTSQRHYMGEDLNTLCLKELQNLEHQLAASLKRIRTRENQLMNKS    150
                   |:|.|:.::|::|||||::|.|||||:|||||.|::|.||:|:||.|.:|
AtFUL          101 RVEVLEKNKRNFMGEDLDSLSLKELQSLEHQLDAAIKSIRSRKNQAMFES    150

OeFUL          151 IAELKKKDKALQDENTLLLKKIKE-EKELSQ---KLQQEQNHDIISSSV-    195
                   |:.|:||||||||.|..||||||| ||:..|   :|.|..|    ||||
AtFUL          151 ISALQKKDKALQDHNNSLLKKIKEREKKTGQQEGQLVQCSN----SSSVL    196

OeFUL          196 -PQ-LLSSGEHGEVE-------GQSQ--SSNAVIPPWMINHLHESNRRG    233
                    || .::|...|.||       |.|.  ..|:::|.||:.....:.
AtFUL          197 LPQYCVTSSRDGFVERVGGENGGASSLTEPNSLLPAWMLRPTTTNE---    242
```

### Additional Materials and Methods

*RNA extraction and first-strand cDNA synthesis*.

Total RNA extraction from leaves was performed using the improved guanidine method (Logemann *et al.*, 1987), and mRNA was isolated using magnetic oligo-dT beads (Dynabeads Oligo $(dT)_{25}$, followed by cDNA synthesis using SuperScript II Reverse Transcriptase (Invitrogen) and Oligo $(dT)_{12–18}$ primers. as previously described (Haberman *et al.*, 2017).

*Expression analysis by quantitative real-time PCR.*

Gene expression was analyzed using ABsolute Blue QPCR SYBR Green ROX Mix (Thermo Scientific) with specific primers (Supporting Information Table S1). Reactions were run on a Rotor-Gene 6000 using a gene encoding Actin 7, OE9A117728 as a housekeeping gene similar to what was previously published (Haberman *et al.*, 2017).

*2021-2022 28$^o$C heat pulses*.

Potted olive 'Barnea' cultivar trees that were at least 2 years old, were used during the winter of 2021-2022 in the following experiment located at the Faculty of Agriculture of the Hebrew University in Rehovot, Israel. The trees were divided into three treatments, and for each treatment, we used 5 trees. In the control treatment, the trees were kept in a net house during the winter. Trees in the 'heat pulse 'treatment were also kept in the same net house during winter, yet were moved four times, once a week, for three hours, starting January 12$^{th}$ 2022, to a controlled temperature room set at 28$^o$C. In one treatment the trees received the heat pulse mid-day, starting at noon. In a different treatment, they received the heat pulse in the evening, starting at 19:00.

*2021-2022 16/10$^o$C followed by different temperature regimes*.

Potted olive 'Barnea' cultivar trees that were at least 2 years old, were used during the winter of 2021-2022 in the following experiment located at the Faculty of Agriculture of the Hebrew University in Rehovot, Israel. Trees from all treatments were moved on October 25$^{th}$, 2021 to nearby controlled-environment 'phytotron' rooms. Here, trees were exposed to day temperatures for 9 hours and night temperatures for 12 hours per calendar day. Changes between day and night temperatures were gradual, spanning 3 h (Sobol *et al.*, 2014). During the first forty-eight days, all phytotron-treated trees were exposed to a 16/10 °C regime. After that, on December 12th, trees were divided into four different temperature regimes for 21 days ((Supporting Information Table S2), until January 2nd, and then moved to a common heated greenhouse (Figure S2). Trees flowered in these conditions and the rate of flowering (*i*) was calculated.

Haberman A., Bakhshian O., Cerezo-Medina S., Paltiel J., Adler C., Ben Ari G., Mercado J.A., Pliego-Alfaro F., Lavee S. & Samach A. (2017) A possible role for FT-encoding genes in interpreting environmental and internal cues affecting olive (Olea europaea L.) flower induction. *Plant Cell and Environment*, **40**, 1263-1280.

Logemann J., Schell J. & Willmitzer L. (1987) Improved method for the isolation of RNA from plant tissues. *Anal Biochem*, **163**, 16-20.

Sobol S., Chayut N., Nave N., Kafle D., Hegele M., Kaminetsky R., Wunsche J.N. & Samach A. (2014) Genetic variation in yield under hot ambient temperatures spotlights a role for cytokinin in protection of developing floral primordia. *Plant Cell and Environment*, **37**, 643-657.
